# Allosteric Mechanisms Underlying Long QT Syndrome Type 2 (LQT2)-Associated Mutations in hERG Channels

**DOI:** 10.64898/2026.04.05.715988

**Authors:** Audrey Deyawe Kongmeneck, Geraldine San Ramon, Brian Delisle, Peter Kekenes-Huskey

## Abstract

Long QT syndrome Type 2 (LQT2) is a genetic disorder caused by missense mutations in the KCNH2 gene that encodes the potassium channel K_V_11.1. Previous studies have shown that most K_V_11.1 missense mutations with loss-of-function phenotypes result from impaired trafficking from the endoplasmic reticulum to the plasma membrane. To investigate the molecular basis of these defects, we used molecular dynamics simulations to analyze two sets of disease-associated missense mutations: those that suppress and those that maintain normal channel trafficking. We focused initially on the conformational and dynamics differences between wild-type and several mutants of K_V_11.1 via molecular dynamics simulations when two K^+^ were placed in the selectivity filter (SF). Our study reveals that missense mutations in the S4 helix allosterically disrupt the selectivity filter, a critical determinant for proper channel trafficking. Trafficking-competent variants largely retained a wild-type selectivity filter structure, whereas trafficking-deficient mutants exhibited pronounced structural perturbations in this region. These findings suggest that certain LQT2-associated missense mutations in KCNH2 impair channel trafficking by compromising the structural integrity of the selectivity filter. We additionally found that second-site variants Y652C in the drug binding vestibule can correct structural defects associated with some mistrafficking variants.

## 2 Intro

### 2.1 Importance of K**_V_**11.1

The K_V_11.1 channel results from the expression of the KCNH2 gene and is mostly expressed in cardiomyocytes, where it participates predominantly late in the cardiac action potential. Kv11.1 slowly activates during membrane depolarization but rapidly inactivates, limiting outward current during the plateau. [1, 2]. K_V_11.1 rapidly recovers from inactivation during the repolarization phase, further driving membrane repolarization after which the channel closes during the cardiac action potential’s resting phase [3].

Numerous loss-of-function missense mutations in the KCNH2 gene are linked to type 2 Long QT syndrome (LQT2) [1]. Long QT syndromes (LQTS) are genetic diseases defined by a prolonged cardiac action potential, visible as an extended QT interval on an electrocardiogram. This prolongation is linked to life-threatening cardiac events including syncope, sudden death, or ventricular tachycardia (Torsade-de-Pointes) and are often triggered in a gene-specific manner [4, 5]. LQT2, which accounts for 30-40% of arrhythmias [6], events are typically induced by sudden sympathetic stimulation during prolonged parasympathetic activity, such as a clock alarm or sudden emotional distress [4]. Because of its critical role, K_V_11.1 is a major target for drug safety, and assessing a drug lead’s potential to affect K_V_11.1 gating is an essential standard step in pharmaceutical development [7, 8]. Therefore, understanding K_V_11.1’s structure and conformational space at a molecular scale is vital for developing functional modulation strategies.

K_V_11.1 is a tetramer, with each subunit comprising a cytoplasmic N-terminal Per-ARNT-Sim (PAS) domain, four transmembrane (transmembrane) helices (S1-S4) forming the voltage-sensing domain (VSD), two transmembrane helices (S5-S6) forming the pore domain (PD), and a cytoplasmic C-terminal cyclic nucleotide-binding domain (CNBD). Its S6 helix and the CNBD are connected by the C_LINKER_, an ensemble of three helices. Functionally, S4 contains highly conserved basic residues that contribute to activation and deactivation, while S6 responds to these signals to configure the highly conserved selectivity filter (SF), which governs K^+^ flux and permeability (Fig. 1).

**Figure 1:**
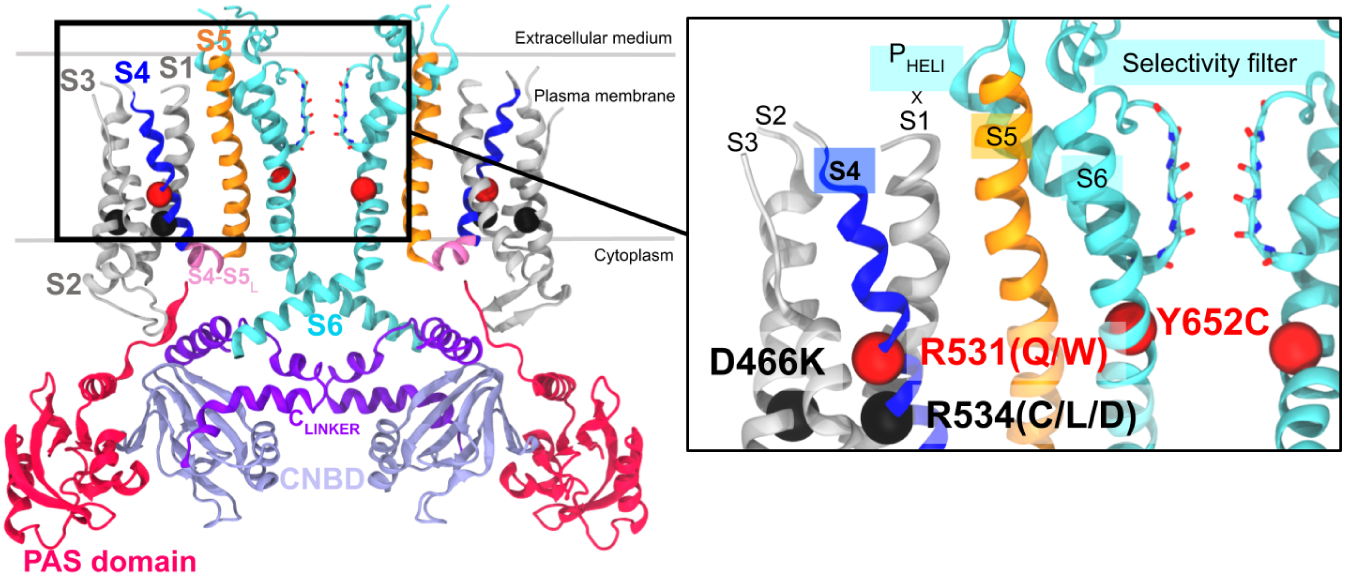
Representations of the K_V_11.1 channel encoded by the KCNH2 gene. Two monomers out of four are shown for clarity. Transmembrane helices S1-S5 are labeled in one monomer of the tetrameric channel. The N- and C-terminal cytosolic PAS (pink) C_LINKER_ (purple) and CNBD (lilac) regions are labeled. The variants considered in this study are rendered in the subpanel relative to the selectivity filter. Non-trafficking variants (Class II) are depicted in black spheres, while non-conducting variants are depicted in red spheres.

### 2.2 Impact of Variants

LQT2-causing KCNH2 variants are classified into four loss-of-function categories [9, 10]: Class I (defective synthesis), Class II (defective trafficking), Class III (defective gating), and Class IV (defective permeation). The majority of LQT2 variants result from Class II missense mutations [11, 12]. Intriguingly, mutations localized to the S4 region can have dramatically different impacts on channel function, ranging from defects in trafficking to shifted voltage dependence of inactivation [13, 14], yet a mechanistic basis for the divergent phenotypes are unknown. As an example, two conserved gating charges, R531 (R4) and R534 (R5), fall into different variant classes due to distinct trafficking outcomes. R4 mutants R531Q and R531W traffic normally (Class III) but exhibit altered gating, with studies showing strongly right-shifted voltage dependence of activation and accelerated activation rates [13, 15–17]. Conversely, R5 mutants R534C and R534L exhibit defective trafficking (Class II). This suggests that the divergent phenotypes observed for sequence-adjacent residues are not merely the result of localized chemical alterations, but reflect a substantial shift in the channel’s global conformational landscape. In our previous structural studies of the KCNH2 PAS domain, we found that some missense mutations induced subtle changes in the domain’s molecular structure and free energy, which we found correlated with trafficking behavior [10]. We proposed that the misfolded structures are subject to retention by the endoplasmic reticulum (ER) quality-control machinery [12, 18]. These observations motivated our hypothesis that S4 variants, even when separated by one helical turn, present different loss of function classes due to unique disruptions of the channel’s conformation ensemble.

### 2.3 Approach

To test our hypothesis, we used MD simulations to predict structural changes in four variants (R531Q, R531W, R534C, R534L). These were compared with control simulations of the wild-type channel and a double charge-reversal mutant, D466K-R534D intended to preserve the wild-type structure. We additionally tested whether the S6 missense mutation Y652C, a variant within the channel’s drug binding vestibule in S6, ‘corrects’ misfolded conformations, based on reports that it restores trafficking for selected Class II mutants [19]. Following the observed effects of S4 variants on the pore structure, we also harnessed MD simulations to investigate whether S4 mutations reshape the structural coupling network linking the VSD, pore domain, and C_LINKER_.

## 3 Materials and methods

The initial coordinates for the molecular dynamics (MD) simulations were derived from the cryo-electron microscopy (Cryo-EM) structure of the K_V_11.1 channel (PDB ID: 5VA1 [20]). Unresolved regions (residues 132–397, 433–448, 511–519, 578–582, and 598–602) were capped as chain termini with amine and carboxylate groups. Single and double missense mutations were introduced using the CHARMM-GUI [21] server based on the wild-type structure. The protein was embedded into a homogeneous 1-palmitoyl-2-oleoyl-sn-glycero-3-phosphocholine (POPC) lipid bilayer using the CHARMM-GUI [21] Membrane Builder [22, 23], for which the top and bottom layer of the membrane comprised 206 and 220 POPC molecules, respectively. The system was solvated with TIP3P [24] water molecules extending 15 °A above and below the membrane. To achieve electrical neutrality and established 160 mM ionic strength, 137 K^+^ and 157 Cl^−^ ions were added. The final assembled systems, comprising approximately 248,000 atoms, were parameterized using the AmberTools tleap utility with the Lipid14 [25] and ff14SB [26] force fields. As illustrated in Fig. 2, two K^+^ ions were initially placed within the selectivity filter (SF) between residues Thr623 and Gly628 consistent with [27].

**Figure 2:**
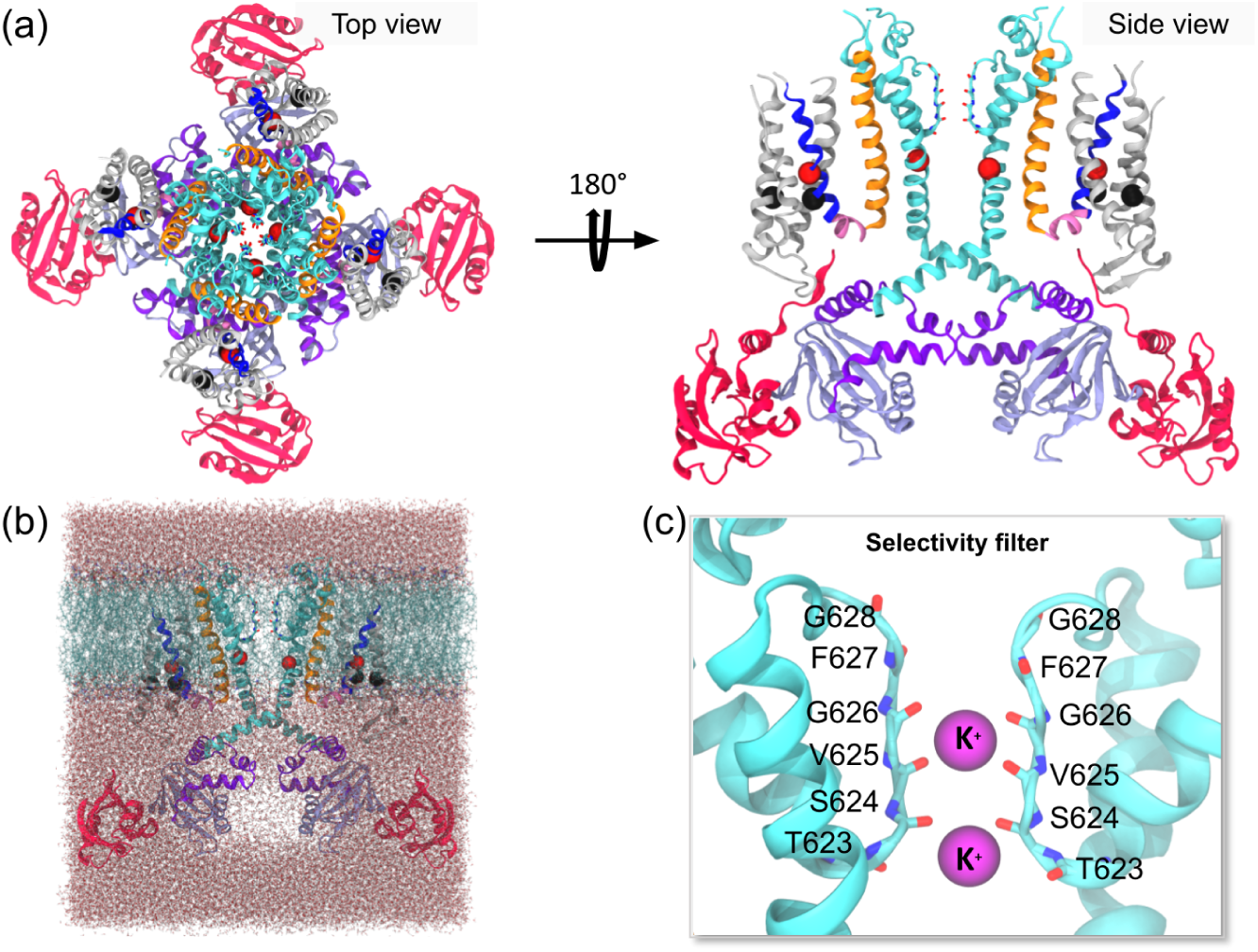
Building Molecular Dynamics systems of K_V_11.1 channel. a) Top-view and side-view of the tetrameric structure of K_V_11.1 channel domains (VSD in gray, S4 in blue, S4-S5_LINKER_ in pink, S5 in orange, pore domain in cyan, C_LINKER_ in purple and CNBD in lilac). Residues that will be mutated according to Class II and Class III missense mutations are depicted in black and red spheres, respectively. b) K_V_11.1 structure embedded into POPC bilayer and two slabs of water molecules. Ions are omitted for clarity. c) Two K^+^ ions were placed in the SF. Only two opposing subunits were shown for clarity.

All simulations were performed using the Amber18 [28]. Energy minimization comprised 5,000 steps (including 250 steps of steepest descent) with gradually decreasing restraints on the lipids and peptide backbone. System heating was conducted in two stages: a 5 ps phase from 0 to 100 K in the NVT ensemble, followed by 100 ps from 100 to 300 K in the NPT ensemble. While the initial minimization utilized the pmemd engine, all subsequent heating, equilibration, and production phases were executed using the GPU-accelerated pmemd.cuda module. For these simulations, the SHAKE algorithm [29] was employed to constrain bonds involving hydrogen atoms, allowing for a 2 fs integration time step. Non-bonded interactions were calculated using a 10 °A cutoff. Temperature was regulated via a Langevin thermostat. During equilibration, heavy atoms of the channel and POPC molecules were initially restrained with a harmonic force constant of 5 kcal/mol. This was followed by two 1 ns equilibration stages at 300 K, where the force constants were sequentially reduced to 3 kcal/mol and 1 kcal/mol. Finally, triplicate production runs of 1 *µ*s each were performed for each system at 300 K without any position restraints. Ions introduced in the system preparation were not subjected to harmonic constraints and tended to diffuse from the SF into the bulk solvent.

MD trajectories were analyzed using a custom pipeline to evaluate the structural and dynamic impact of the mutations. This included monitoring the Root Mean Square Deviation (RMSD) and Root Mean Square Fluctuation (RMSF) of C*_α_* atoms, time-dependent pore profiles and the torsion angles of the SF backbone, sampled every 2 ns. Intersubunit and interhelical interactions were additionally characterized between residues critical to activation and inactivation mechanisms, those forming binding sites for pharmacological correctors (e.g., dofetilide and E-4031) [30–32], and those associated with Class II missense mutations [11, 33]. This comprehensive analysis quantified side-chain interactions, pore geometry, and interdomain correlations.

## 4 Results

Variants at R531 and R534 were specifically selected from the S4 helix due to their distinct trafficking phenotypes despite their sequence adjacency (Fig. 2). We additionally included a Y652C missense mutation to test if it could restore interactions potentially disrupted by the S4 variants, and a D466K-R534D double mutant as a control. All cases were subjected to triplicate 1 *µ*s simulations of both the wild-type (wild-type). We monitored these simulations for global conformational changes and observed that the overall channel structure did not change appreciably relative to wild-type, regardless of the severity of the associated trafficking or gating defects. This structural stability was confirmed by similar root mean squared deviations (RMSD) values for C*_α_* atoms (Fig. S1) and root mean squared fluctuations (RMSF) profiles that revealed consistently low mobility within the transmembrane region (residues 398-687) compared to the cytoplasmic domains (PAS: 1-131; C_LINKER_: 688-741; CNBD: 742-863). These analyses of the overall channel structure and dynamics indicate that the protein is not significantly impacted by the missense mutations at the microsecond timescales accessible to our simulations.

### 4.1 Pore and selectivity filter structural states

While the LQT2-associated variants investigated in this study are localized to the S4 segment, their pathological phenotypes manifest as impaired channel function, often linked to the pore domain and SF regions which are enriched in loss-of-function variants [11, 33].

Given the selectivity filter’s central role in ion conduction, C-type inactivation, and pharmacological interactions, we therefore focused our structural analyses on this region. Consequently, we investigated whether S4 missense mutations allosterically impact the pore structure.

#### 4.1.1 Pore size

To quantify the openness of the pore region, we used HOLE [34] to iteratively measure pore radii along the conduction pathway in 0.125 °A increments, from MD coordinate data sampled every 20 ns (Fig. 3). Our 1 *µ*s MD simulations indicated that all K_V_11.1 variants maintained a pore structure similar to wild-type at the SF level, where the average radius remained below the 1.33 °A ionic radius of K^+^, confirming the desolvation of K^+^ ions during diffusion (Fig. 3a,b) [35, 36]. We next assessed the conducting pathway using Connolly surfaces and observed that while there was no uniform constriction, the profiles of the non-trafficking variants suggested a loss of fourfold symmetry (Fig. 3c,d). To resolve the basis of this broken symmetry, we quantified inter-carbonyl distances between opposite V625 residues and found that these distances increased in non-trafficking mutants (Fig. S4a). These assessments of the pore structure and subunit-specific rearrangements reveal that S4 mutations induce a loss of fourfold symmetry and increased inter-carbonyl distances at the selectivity filter. To identify the structural origin of these observed changes, we compared pore radii profiles with Cryo-EM structures solved under high- and low-[K^+^] conditions. We first partitioned the R534C and R534L datasets into two groups based on the V625 main chain conformation, which exhibits distinct carbonyl peptide bond orientations directed inside or outside the pore (Fig. 3c,d). We found that the R534 missense mutations appear to shift the SF carbonyls downwards along the Z axis towards the inner membrane, explaining the offset between the wild-type curve (black) and those of the mutants R534C (pink) and R534L (orange) (Fig. 3a,c). In the specific case of R534L, the orientation of V625 did not appreciably change the pore radius (≈ 0.25 °A). This contrasts with K_V_11.1 Cryo-EM pore radii (Fig. 3e-f), where structures show significant differences at the SF level due to distinct V625 orientations: V625∗in for high [K^+^] and V625∗out for low [K^+^]. These results confirm a reduced solvent-accessible surface of the pore when the SF is inactivated, although the magnitude of the radius change in R534L/C is smaller than observed in Cryo-EM structures. This difference occurs because the V625∗out conformation manifests in only one or two subunits rather than all four (Fig. 3b,d).

**Figure 3:**
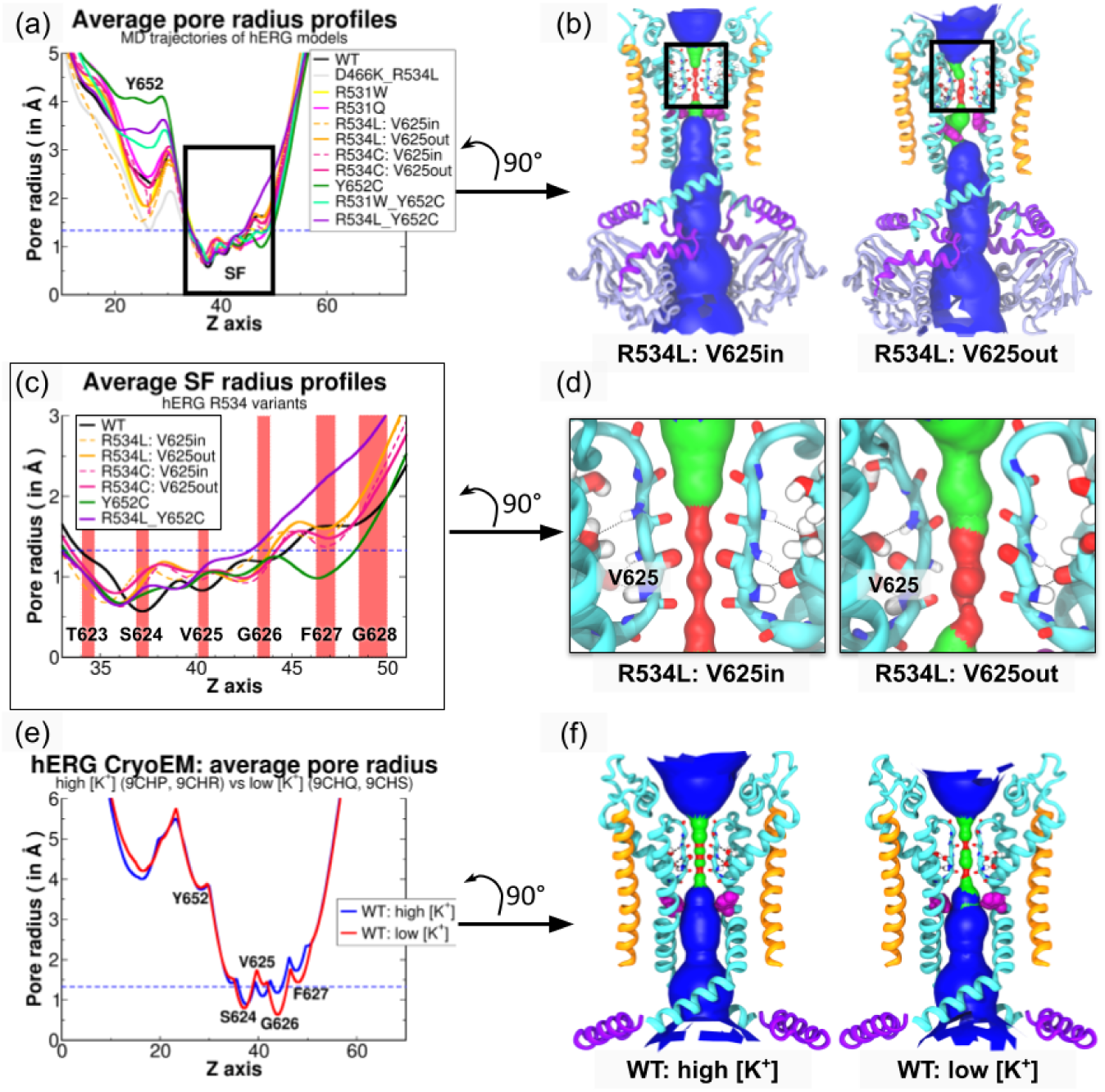
Average pore size of K_V_11.1 channel variants over triplicate MD simulations. Panels (a), (c) and (e) show average radius profiles of the entire pore domain in all K_V_11.1 variants, the SF in R534 mutants, and the Cryo-EM structures of wild-type K_V_11.1 in high and low [K^+^] environments [37], respectively. In panel (c), red areas represent positions occupied by carbonyl oxygen atoms of each SF residue throughout the wild-type K_V_11.1 MD trajectory. Panels (b), (d) and (f) depict the average accessible solvent surface of the K_V_11.1 R534L variant according to the two different V625 carbonyl peptide bond orientations (relative to the conduction pathway center) and the wild-type K_V_11.1. K^+^ ions in the pore are ommitted for clarity. Pore surfaces are colored according to calculated average pore radii. Values below 1.33 °A (K^+^ ionic radius) are colored red. Values ranging from 1.33 °A to 3.6 °A (K^+^ hydrodynamic radius) are colored green. Values above 3.6 °A are colored blue.

We next examined whether R534 trafficking-deficient mutations impacted the structural organization of the selectivity filter (SF). Pore size was accordingly quantified using distances between opposing residues across the pore. Our measurements of the average G626 C*_α_* distances from opposite subunits in the K_V_11.1 models (Fig. S4b) confirmed that the pores likely adopted an active state, which is characterized by a symmetric conductive conformation. These results align with free energy calculations by Li et al. [27], for which PMF profiles based on opposing G626 C*_α_* distances were computed. This literature suggests that 8 °A corresponds to the symmetric conductive state, whereas 5-7 °A corresponds to asymmetric stable inactive states, similar to the 5.5 °A energy well observed in the non-conductive state of the homolog K^+^ channel KcsA. Hence, we extended our analysis of SF symmetry by measuring distances between opposing backbone carbonyls that could interact with K^+^. We found that carbonyl distances associated with V625 (Fig. S4a) in R534C and R534L single mutants increased significantly compared to wild-type and trafficking-competent variants.

#### 4.1.2 SF backbone conformational changes and solvent accessibility

We next evaluated backbone torsions to better characterize the conformational changes exhibited by V625. We further analyzed backbone conformational changes using Ramachandran plots of selectivity filter (SF) residues, to assess torsion angles in our MD simulations (Fig. 4) We also included P_HELIX_ residue S620 as a control for V625 switching between in and out conformations. The results revealed significant differences in the regions occupied by the backbone angles of V625 and G626. Non-trafficking mutants populated distinct conformational regions relative to wild-type, whereas trafficking-corrected mutants recovered WT-like backbone conformations. Specifically, the Ψ values of V625 and G626 spread toward more positive values (+70^◦^) in non-trafficking variants compared to non-conducting, wild-type, and double mutant channels. Although V625 Φ values remained unchanged, those of G626 spread toward more negative values in non-trafficking variants versus wild-type and trafficking-competent ones. While the wild-type channel maintains a dynamic equilibrium between conductive states, the non-trafficking variants appear to be allosterically driven into a structural entrenchment, where the selectivity filter is asymmetrically trapped in a non-native, solvent-inaccessible configuration. Importantly, these data suggest that observed changes in V625 opposing carbonyl distances in non-trafficking K_V_11.1 variants arise from altered SF backbone torsion angles.

**Figure 4:**
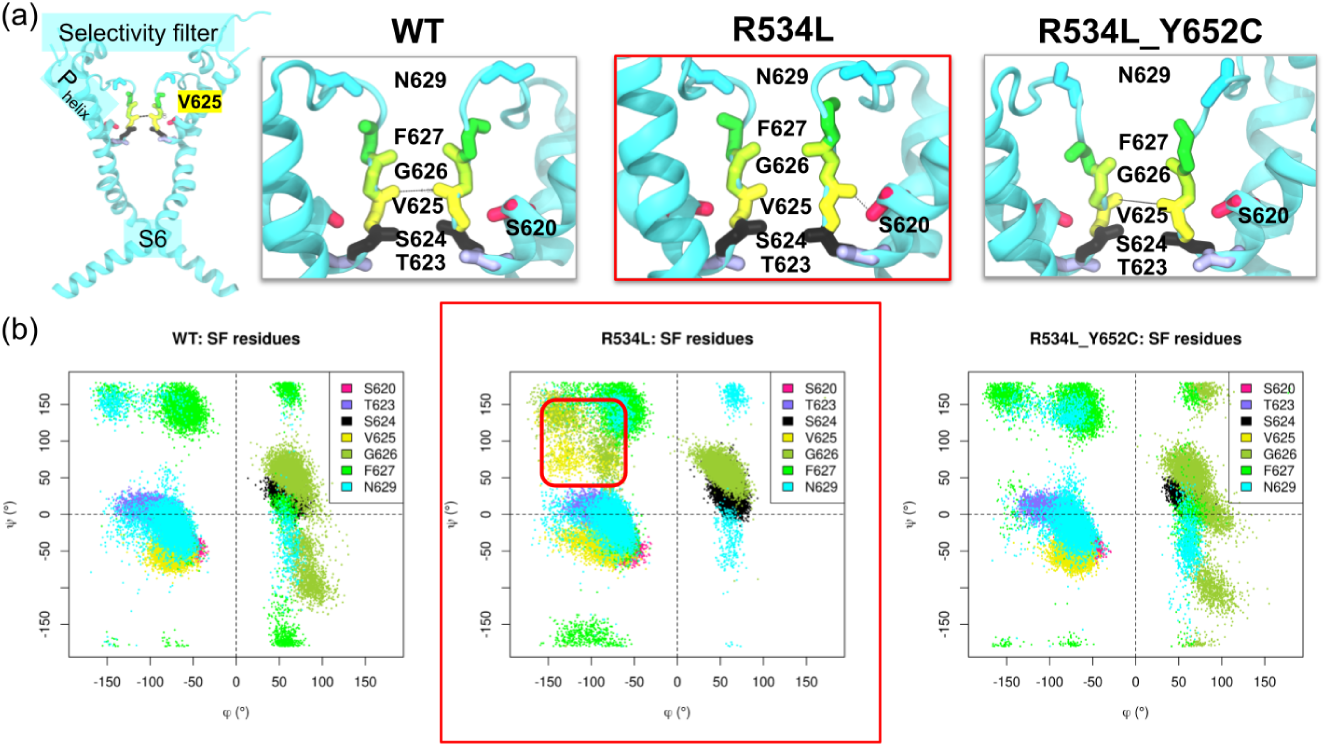
Ramachandran plots of selectivity filter residues in K_V_11.1 variants. (a) Cartoon representation of K_V_11.1 pore domain in the wild-type (left) and enlarged views of K_V_11.1 selectivity filters in the wild-type, the single mutant R534L (framed in red) and the double mutant R534L-Y652C. In each panel, the pore domain is colored in cyan. The sidechain of P_HELIX_ residue S620 (pink) and the main chain of SF residues T623 (lilac), S624 (black), V625 (yellow), G626 (lime green), F627 (green), and N629 (cyan) are represented in sticks. (b) Ramachandran plots of the Φ/Ψ torsion angle values (in degrees) of P_HELIX_ S620 and SF residues T623, S624, V625, G626, F627 and N629 in the MD trajectories of K_V_11.1 wild-type (left), R534L (middle, framed in red), and R534L-Y652C (right) variants. The area occupied by the Φ/Ψ values for V625 and G626 in the MD trajectory of the non-trafficking K_V_11.1 mutant R534L is framed in red.

#### 4.1.3 Trafficking-dependent selectivity filter interactions

We next examined specific interactions associated with distinct SF states (Fig. 5). According to findings from [37], hydrogen bonds between the S620 side chain and main chain oxygens of Y616 and V625 reportedly stabilize partially- and fully-inactivated K_V_11.1 SF states, respectively (Fig. 5b). Conversely, the activated state is characterized by hydrogen bonds between the S620 main chain oxygen and the main chain nitrogens of V625 and G626, together with a hydrogen bond between the S620 hydroxyl oxygen and the F627 main chain nitrogen (Fig. 3e-f). We quantified this deformation using the H-bond interaction probability, calculated as a portion of the integrated probability density function of the distance between H-bond donor and acceptor atoms, such as the V625 backbone oxygen and S620 side chain oxygen (Fig. 5c). Our analysis revealed significantly larger binding areas for R534C and R534L variants (Fig. 5d), suggesting V625 engaged in a hydrogen bond with S620 that may trigger an asymmetric, inactive SF conformation. Interestingly, we also found the S620-Y616 interaction in our non-trafficking R534L and R534C models, but not in the wild-type or other K_V_11.1 variants (Fig. 5b,d). These results, which are consistent with structural changes reported for low-[K^+^] environments [37], suggest that the increased distance between V625 carbonyl oxygens in non-trafficking R534 mutants is stabilized by a hydrogen bond between the V625 main chain and the S620 side chain. (Fig. 5a,b)

**Figure 5:**
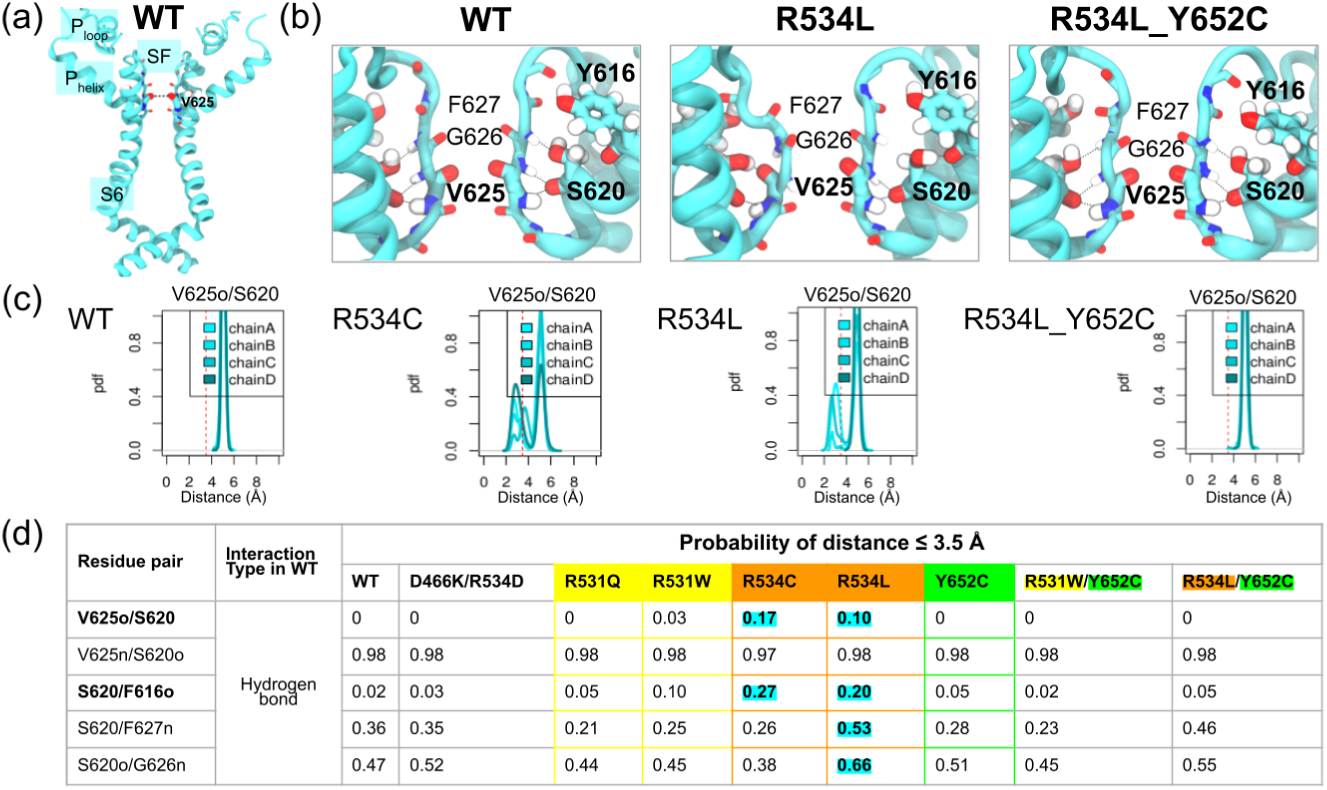
Identifying K_V_11.1 trafficking-dependent interaction in the pore domain. (a) Cartoon representation of the K_V_11.1 pore domain in the wild-type (WT). (b) Enlarged views of selectivity filters of K_V_11.1 wild-type (left), single mutant R534L (middle), and double mutant R534L-Y652C (right). (c) Probability density functions of the interatomic distance between the V625 backbone oxygen and S620 side chain oxygen for every subunit (chains A-D) of wild-type and variants R534C, R534L, and R534-Y652C. The 3.5 °A threshold characterizing hydrogen bonds is represented by a vertical dashed red line. (d) Table reporting the percentage of the area under the probability density curves falling below 3.5 °A obtained in panel (b) throughout the MD trajectories of all K_V_11.1 variants.

Collectively, the Φ/Ψ angle changes (Fig. 4, Fig. S6) sufficiently impacted the backbone to allow S620-V625 hydrogen bond formation. We speculate that G626 backbone perturbation might shift the equilibrium from the symmetric conductive state toward the asymmetrical non-conductive state described by Li et al., which is defined by reduced solvent accessibility. These results coincided with structural changes associated with low external [K^+^] in the K_V_11.1 selectivity filter (SF) [37]. Indeed, this study confirmed that the S620 side chain binds F627 backbone nitrogens in its active conformation as shown previous in [38], switches to the P_HELIX_ residue Y616 backbone oxygen in an intermediate state, and finally to the V625 backbone oxygen in the fully inactive state as the latter flips its main chain carbonyl. However, the S620-F627 H-bond was maintained in our R534 single mutants (Fig. 5d), unlike in the low [K^+^] structure where it was abolished. This difference likely occurred because the V625 backbone flip did not occur in all subunits.

### 4.2 S4 variants impact TM position and packing

Because the trafficking defect originates from missense mutations in the VSD, we investigated how electrostatic interactions involving S4 gating charges were altered in our models before examining how these perturbations propagated to the pore architecture. Proceeding this way allowed for the identification of the structural origin of the trafficking defect associated with R534 missense mutations. Although K_V_11.1 S4 presented five of seven strongly conserved gating charges observed among K_V_ channels (Fig. 6a), K525(K2), R528(R3), R531(R4), R534(R5), and R537(R6), along with three K_V_-conserved countercharges, D456(D1), D466(D2) from S2, and D501(D3) from S3, its VSD is unique. Specifically, this VSD contains an additional positive S4 charge, K538(K6’), and three extra countercharges: D411 from S1, D460(D1’) from S2, and D509(D3’) on S3. Therefore, we specifically examined its electrostatic network to determine a structural basis for trafficking-deficient mutations.

**Figure 6:**
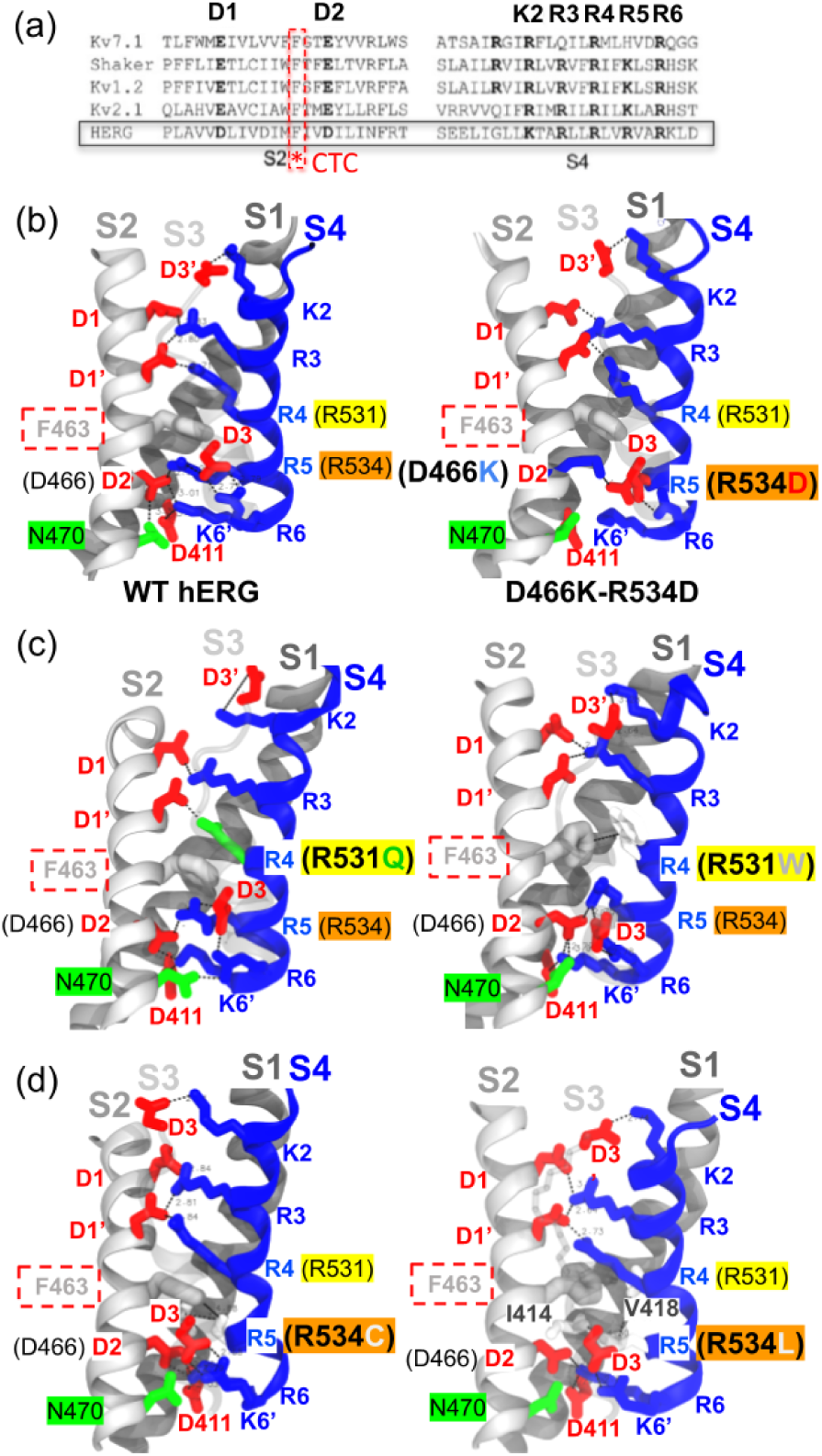
Identifying K_V_11.1 trafficking-dependent interactions in the pore. (a) Sequence alignment of transmembrane helices S2 and S4 in Voltage-Gated K^+^Channel homologs K_V_7.1, Shaker, K_V_1.2, K_V_2.1, and K_V_11.1 (black frame). The conserved charge transfer center (CTC) F463 is framed by a red dashed line. (b) Structural mapping of conserved S2 (solid gray), S3 (transparent gray), and S4 (blue) residues and interactions in the wild-type model (left) and WT-like double mutant D466K-R534D (right). Residues are shown as sticks; acidic residues are red, basic residues are blue. The CTC is colored gray. (c) and (d) show the structural mapping of S2-S4 interactions in K_V_11.1 R531 (R531Q, R531W) and R534 (R534C, R534L) variants, respectively. Color coding matches panel (b).

#### 4.2.1 Assessment of S4 position and contacts

In experimental structures of wild-type K_V_11.1 transmembrane/cytoplasmic regions (PDB: 5VA1) and recent transmembrane structures in high (PDB: 9CHP, 9CHR) and low (PDB: 9CHQ, 9CHS) [K^+^] environments, salt bridges were identified between pairs K2-D3’, R3-D1, R3-D1’, R4-D1’, R5-D2, R5-D3, and R6-D3 (Fig. S7a). Additionally, the charge transfer center (CTC) formed cation-*π* interactions with both R4 and R5, while D2 engaged in a hydrogen bond with N470 in both wild-type and high [K^+^] transmembrane structures. Interestingly, while the wild-type K_V_11.1 transmembrane/cytoplasmic structure showed an additional K6’-D2 salt bridge, the K_V_11.1 transmembrane structures showed K6’ binding D411 on S1 instead. These experimental structures suggest that the primary difference between R4 and R5 was that the R4 side chain was positioned above the CTC, whereas R5 was located below the CTC (Fig. S7a), stabilizing the VSD in its activated state. Meanwhile, R5 remained below the CTC alongside R6, K6’, S1 residue D411, S2 residue N470, D2, and D3. Together, these residues formed the occluded site, shielded from the extracellular solvent by the CTC side chain. Taken together, the observations of K_V_11.1 cryo-EM structures suggest that S4 gating charges remaining occluded from the extracellular medium contribute to K_V_11.1 function through electrostatic stabilization rather than voltage-driven translocation. This stabilization shifts the thermodynamic equilibrium, biasing the channel toward conformational ensembles associated with either the activated or inactivated states.

To evaluate this hypothesis, we quantified the dynamics of these electrostatic interactions throughout the MD trajectories obtained for our K_V_11.1 models to assess differences in the VSD salt-bridge network. Toward this end, we monitored interatomic distances between charge moieties of S4 gating charges and their VSD countercharges on S1, S2, and S3 using MD trajectories of wild-type and variant models. We represented electrostatic interactions as integrated probability densities of interatomic distances falling below 3.5 °A throughout each trajectory (Fig. S7b). These data indicated that most specific interactions found in K_V_11.1 cryo-EM structures (Fig. S7a) were maintained in the wild-type model trajectory (Fig. 6b). In R531 single mutant trajectories, the R4-D1’ and R5-D2 salt bridges along with R4-CTC cation-*π* interaction were impaired, but only R5-D2 was restored to WT-like values in R531W-Y652C, indicating that R4-D1’ and R4-CTC interactions were not restored by Y652C (Fig. S7b). In comparison, the R534 single mutants drastically altered interaction probabilities for the K2-D3’ salt bridge and R5-CTC cation-*π* interaction while completely abolishing R5-D2 and R5-D3 (Fig. 6d), thereby disrupting the electrostatic network of the occluded S4 binding site (Fig. S7b). Instead, R534 single mutants established hydrophobic contacts with S1 residues I414 and V418. We believe that the addition of the Y652C missense mutation might restore the K2-D3’ salt bridge and R5-CTC cation-*π* interaction to WT-like values through a coupling mechanism between the VSD and the pore, but not through the restoration of the occluded site as the electrostatic interactions involving R5, as R5-D2 and R5-D3 both remained impaired in R534-Y652C MD trajectories.

Interestingly, we found that D466-N470 hydrogen bond probabilities were lower in non-trafficking mutants compared to other variants. Given that these residues are associated with non-trafficking phenotypes [39], this suggests that their hydrogen bond might play a crucial role in VSD gating charge network integrity. In further support of this conjecture, we found that the probabilities of R6 and K6’ forming salt bridges with occluded site residues D3 and D2, respectively, were significantly higher than wild-type in both R531 and R534 single mutants (Fig. S7b). The potential importance of this observation is that R6 and K6’ are known to participate in activation and deactivation voltage dependence [14, 16], in part through K6’ interacting with S1 via residue D411 [40]. However, K6’-D411 salt-bridge showed similar or higher probabilities than K6’-D2 in most MD trajectories (Fig. S7b). Overall, because S4 gating-charge and VSD counter-charge missense mutations modulate activation, deactivation, and inactivation-related metrics, including voltage-dependence and recovery, these data supported a model where VSD perturbations reshape pore conformational ensembles relevant to inactivation [13–15].

#### 4.2.2 S4-pore coupling: S4-S5_LINKER_ torsion angle measurement

To identify other potential mechanisms of S4-pore coupling, we analyzed the time-dependent torsion angle between S4 and S5 helices relative to the S4-S5_LINKER_ axis (Fig. S5). We found that changes in this dihedral angle (which is decreased only in R534 single mutants compared to the wild-type) are mildly correlated with the trafficking-competent phenotypes of the rescued double mutants. Hence, we speculate that the relative packing of the S4 and S5 segments in these variants presents a conformation compatible with ER export rather than retention. Altogether, these observations is in agreement with the idea that alterations in steady-state inactivation voltage-dependence, rather than gross defects in folding or kinetics, may be a recurring feature of correctable trafficking-deficient K_V_11.1 variants.

### 4.3 Long-range allosteric coupling revealed by correlation matrices

#### 4.3.1 K_V_11.1 PD-C_LINKER_ coupling network: variant-independent interactions

Given the pore differences highlighted above, we sought to determine a plausible molecular mechanism linking local VSD impacts to allosteric effects on the pore structures. We highlight a pore domain (PD) interaction network comprising a bundle of three intrasubunit interactions: hydrophobic contacts S5-SF (F557-T623) and SF-S6 (T623-Y652), and C_LINKER_–C_LINKER_ electrostatic interactions R696-D727^∗^ and P721-Y700^∗^ (Fig. S8a). We first measured the distances between the sidechains of each residue pair in the hERG transmembrane/cytoplasmic Cryo-EM structure and evaluated interaction probabilities for this region based on integrated distance density functions (Fig. S8b) falling below 5 °A for hydrophobic contacts and 3.5 °A for electrostatic interactions throughout MD simulations. Despite being closer to Y652 aromatic ring than the one of F557 in the experimental structure, the interaction probability of T623’s methyl group to bind F557 is around ≈ 0.20, versus ≈ 0.10 to bind Y652 in all MD simulations. Similarily, the guanidium group of R696 was located closer to the carboxylic group of D727 than the D767 one in the experimental structure, yet R696 turned out to interact with both aspartic acids at probabilities around ≈ 0.50 each in all MD simulations. As expected by their distance in the Cryo-EM structure, the hydrogen bond between P721 and Y700 remained equally present in all MD simulations, with probabilities lying around ≈ 0.15. Our data indicate that the PD interactions noted above are largely preserved across all variants we considered. Interestingly, several residues involved in this state-independent bundle are associated with known LQT2 Class II missense mutations (T623I, R696C, R696P, P721L) [11, 33], while the F557L mutant was shown to impair K_V_11.1 interaction with pharmacological correctors [32, 41]. While these PD interactions may be important for the intrinsic stability of the channel’s tetrameric structure, could they contribute to S4 variants’ phenotypes ?.

To connect the observed local structural alterations with global channel dynamics, we computed Pearson correlation matrices across a set of time-dependent distances, including the pore-C_LINKER_ interaction network, the SF residues interactions and the S4 gating-charge interactions along with the dihedral angle between S4 and S5 helix axes (Fig. S5) and report values with *p*-values below 0.05. We calculated these for gating charge interactions (Fig. 6), S4-S5 torsion angles (Fig. S5), SF residue interactions (Fig. 5), and the variant-independent pore domain interaction bundle (Fig. S8). This was done in order to reveal how residue interactions, not just positions, may be allosterically impacted by S4 variants. We consistently observed variant-independent correlations between SF-S5 (T623-F557), SF-S6 (T623-Y652), intersubunit C_LINKER_–C_LINKER_ (R696-D727^∗^ and P721-Y700^∗^), and the intrasubunit C_LINKER_–CNBD interaction R696-D767, suggesting a conserved mechanical framework across all variants. In contrast, correlations involving S4 interaction K525(K2)-D509(D3’) and the non-trafficking specific hydrogen bond V625o-S620 were selectively lost in non-trafficking mutants (whose respective data were merged into one matrix as their coefficients present similar values in Fig. 7b). Similarly, correlations involving intersubunit C_LINKER_ interaction P721-Y700^∗^ and V625o-S620 are strongly anti-correlated in most variants except the non-trafficking ones, meaning that P721 gets closer to Y700 on another subunit when V625 gets farther from S620 in the trafficking competent variants. Despite being absent in trafficking-deficient variants, these long-range couplings were preserved in R531 single mutants (whose data were merged into one correlation matrix in Fig. S9a) and were restored or reorganized in the trafficking-corrected variant R534L-Y652C (Fig. S9b) as well as in the control variants R531W-Y652C and Y652C, whose data were merged into one correlation matrices as they were showing similar correlation coefficients (Fig. S10a). Together, these results reveal a cross-talk between S4 configurations, SF structural states, and C_LINKER_ dynamics, providing a mechanistic basis for how specific missense mutations might disrupt or restore K_V_11.1 channel trafficking.

**Figure 7:**
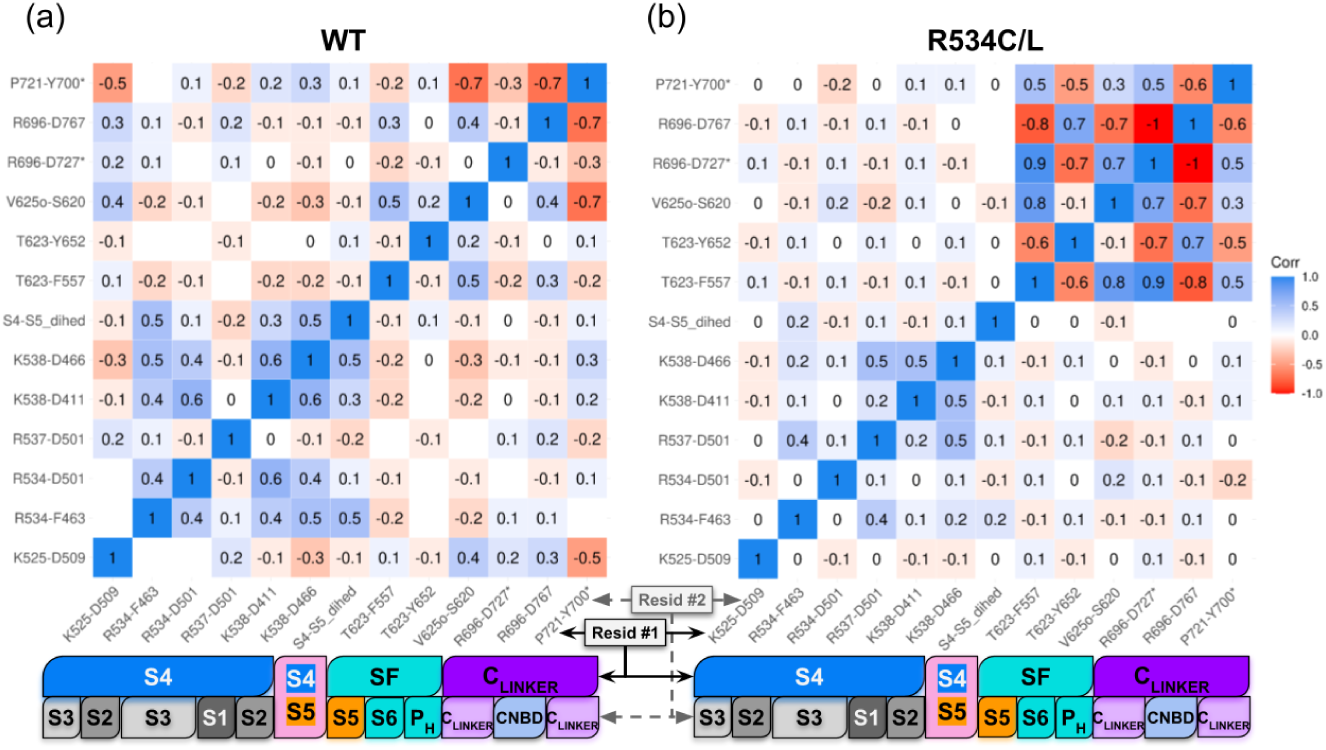
Correlation matrices of pore domain residue interhelical interactions in wild-type and non trafficking K_V_11.1 variants. (a) Pear-son coefficient heatmap of interatomic distances of pore domain residue pairs in MD trajectories of (a) wild-type K_V_11.1 and (b) K_V_11.1 non-trafficking mutants R534C and R534L. For each residue pair, the Pearson coefficient was calculated with p ¡ 0.05; blank areas represent non-significant values. The emergence of high-correlation clusters (red/blue) in the non-trafficking variants reveals an abnormal allosteric pathway where the selectivity filter becomes mechanically locked to the cytoplasmic C_LINKER_ assembly. For both panels, two horizontal bars are colored according to the region of each residue in the corresponding residue pairs.

#### 4.3.2 Differential impact of R4 versus R5 missense mutations on K_V_11.1 trafficking

Given the apparent structural differences in the VSD distal from the S4 mutations despite largely preserved VSD interactions, we propose that the distinct trafficking phenotypes observed for R531 and R534 arise from the selective loss of specific coupling pathways involving R534. For instance, we found for wild-type channels (Fig. 7a) that S4-S5 dihedral motions that were positively correlated with the salt bridge K538-D466 as well as with the cation-*π* interaction between R534 and the CTC residue F463. In other words, as the the S4-S5 dihedral angle increases, the distances K538-D466 and R534-F463 lengthen, even while remaining within typical hydrogen-bond and cation-*π* ranges, respectively; conversely as the dihedral angle decreases, these interactions trend toward shorter values. These correlations suggest a coordinated coupling between S4 positioning and the pore-lining helices. The correlation patterns are largely preserved for the R4 variants R531Q and R531W (see Fig. S9a). This indicates that R4 missense mutations might perturb gating without destabilizing the conformational space that would allow for forward trafficking. In contrast, these correlations were lost in R5 non-trafficking variants (Fig. 7b), suggesting that the trafficking defect induced by R5 missense mutations are connected to the disruption of the VSD-pore coupling network rather than from a local destabilization of the voltage sensor itself. Intriguingly, our double-mutant variant D466K-R534D (Fig. S10b) restored positive correlations between S4-S5 dihedral motions and the mutant, resulting in an anion-*π* interaction R534D-F463, in addition to those between S4-S5 dihedral motions and alternative salt bridges R537-D501 and K538-D411, revealing that the trafficking defect induced by R5 missense mutations originates from a disruption of the VSD4-pore coupling network rather than from a local destabilization of the voltage sensor itself.

The loss of S4-S5 dihedral motion reported for the R5 missense mutations was accompanied by a pronounced reorganization of long-range interactions linking the selectivity filter (SF) to the C_LINKER_. The reorganization gave rise to a rigidified pore architecture evidenced by strong positive/negative correlations (Fig. 7b). Namely, in non-trafficking R5 variants, SF-associated interactions inclusive of T623-F557, T623-Y652, and V625o-S620 became strongly correlated or anti-correlated with intersubunit C_LINKER_ interactions involving R696-D727^∗^, R696-D767, and P721-Y700^∗^ (Fig. 7b). In contrast, these interaction pairs remained largely uncorrelated in the wild-type (Fig. 7a) and non-conductive variants, indicating that the SF and C_LINKER_ are normally dynamically decoupled (Fig. S9a). Further, this abnormal coupling was absent in trafficking-corrected variants (Fig. S9b-Fig. S10). The emergence of strong SF-C_LINKER_ correlations in non-trafficking mutants therefore could reflect a loss of conformational plasticity, whereby SF inactivation-associated states became mechanically locked to the C-terminal tetrameric interface. Together with the observed probabilities of SF interactions associated with inactivated states (Fig. 5d), these data support a model in which R5 missense mutations stabilize a non-native, rigidified pore ensemble characterized by excessive intersubunit C_LINKER_ coupling. We speculate that such an ensemble could engage ER quality-control machinery [12, 18, 42] that would lead to the channel’s retention and non-trafficking phenotype. These correlation patterns indicate that trafficking-deficient variants exhibit stronger coupling between fluctuations in the VSD, pore domain, and C_LINKER_, consistent with a rigidified conformational network and reduced structural plasticity.

## 5 Discussion

### 5.1 Summary

Loss-of-function missense mutations associated with LQT2 predominantly arise from impaired K_V_11.1 trafficking rather than permeation defects [11, 12]. We investigated LQT2-linked S4 missense mutations segregating into distinct functional classes: Class III variants (R531W, R531Q), which traffic but exhibit altered gating, and Class II variants (R534C, R534L), which are ER-retained. Although located just one turn apart on the same VSD helix, these mutations induce class-specific perturbations that extend over 30 °A toward both the selectivity filter and the cytoplasmic linker.

Our simulations revealed that missense mutations belonging to the same functional class stabilize similar conformational ensembles, whereas those from different classes shift the channel toward distinct, non-overlapping regions of conformational space. Rather than inducing isolated local defects, Class II missense mutations promote the emergence of a non-native allosteric ensemble characterized by altered coupling between the VSD, the SF, and the C_LINKER_. This abnormal allosteric rigidification of the VSD, SF, and C_LINKER_ likely restricts the conformational plasticity required for proper maturation. In contrast, Class III missense mutations primarily perturb voltage-dependent gating while preserving the structural plasticity that appears to be required for forward trafficking.

A central component of this allosteric ensemble is the conformational plasticity of the SF. Throughout the MD trajectories of K_V_11.1 R534 single mutants, the SF dynamically interconverted between conductive and non-conductive states, a process critically influenced by the orientation of the V625 backbone carbonyls (Fig. 3). The flipping of these carbonyls away from the pore axis, facilitated by interactions with P_HELIX_ residues Y616 and S620, converted the filter into a solvent-inaccessible, non-conductive configuration associated with K_V_11.1’s characteristic rapid inactivation (Fig. 5). Previous simulations have shown that this flipped state of V625 prevents K^+^ permeation by narrowing the filter and introducing an electrostatic barrier in a low [K^+^] environment [37]. While such transitions are intrinsic to normal gating, our results indicated that the Class II missense mutations investigated here biased the conformational ensemble toward these non-conductive states and enhanced their coupling to distal regions of the channel.

In addition, our proposed SF-C_LINKER_ coupling axis aligns with experimental datasets showing that substitutions along this pathway produce Class II variants with distinct correctability profiles. Several PD/C-terminal variants along the SF-C_LINKER_ continuum (e.g., V625E, G626D/V, R696P, P721L) are refractory to low temperature or E-4031 rescue, whereas others (e.g., G626S, R696C) remain correctable. Furthermore, the VSD CTC substitution F463L severely impairs K_V_11.1 trafficking [43], while the CNBD variant D767Y is a Class III mutation [33]. Hence, the pore-to-C_LINKER_/CNBD axis may play a critical allosteric role in governing trafficking efficiency.

Beyond R5 mutations, this mechanistic framework extends to other Class II LQT2 variants along the same allosteric axis. For instance, the N470D mutation in the VSD occluded site (Fig. 6b-d, Fig. S7a) likely alters the S4 electrostatic environment, restricting flexibility and biasing the channel toward a non-native ensemble. Consistent with disrupted long-range coupling, N470D is correctable by temperature or E-4031, but not by Y652C [19, 44]. Elsewhere in the channel, our correlation analysis identifies R696 intersubunit electrostatic interactions as dynamically coupled to SF rearrangements in non-trafficking variants. Based on this, we speculate that substituting R696 with cysteine (Class II variant R696C [33]) might impair these interactions, thereby stabilizing aberrant SF–C_LINKER_ coupling and rigidifying the cytoplasmic assembly.

### 5.2 An Alternative hypothesis

Although trafficking-deficient K_V_11.1 variants are often broadly described as misfolded [11, 12, 33], which functional conformations are selectively destabilized remains unclear. An alternative perspective is that trafficking deficiency stems not from an inability to reach activated or deactivated states, but from disruption of the conformational pathway accompanying C-type pore inactivation. We offer this interpretation based on previous experimental and structural studies focusing on S4’s role in K_V_11.1 gating and trafficking. For instance, missense mutations affecting K_V_11.1 S4 gating charges and their counter-charges, including D456Y [11], K525A, R528A [13], and F463L [43], lead to trafficking-deficient phenotypes. Interestingly, the CTC residue F463 (Fig. 6a), a highly conserved S2 residue whose Shaker equivalent (F290) regulates the fast component of S4 gating charge displacement [45], was found to govern the passage of the fourth gating charge across the membrane [46].

In K_V_11.1, only the first three gating charges (K2, R3, R4) move across the electric field generated by membrane polarization changes, as shown in the VSD of our K_V_11.1 models (Fig. 6b-d). As such, they mediate the motion of S4 across the membrane plane during the voltage-dependent activation and deactivation mechanisms [14]. The third gating charge, R4, was found to play a pivotal role in the activation and deactivation mechanisms, as well as in K_V_11.1 gating charge movement [13–16]. Residues R5, R6, and K6’ reportedly remain inaccessible from the extracellular milieu under all tested conditions, while D2 becomes accessible in the deactivated state [14], defining a state-dependent occluded site within the VSD. In K_V_11.1, VSD activation (linked to a closed pore) toward the fully activated state (linked to an open pore) is rapidly followed by C-type inactivation, a pore closure mechanism involving SF rearrangements that render the channel non-conductive despite a depolarized VSD [3]. Despite their lack of transmembrane movement, substitutions at these positions markedly alter steady-state inactivation voltage dependence, shifting it toward less (R534A, R537A) or more (K538A) hyperpolarized potentials compared to wild-type [13]. Moreover, R537C significantly decreases the inactivation rate compared to wild-type, whereas K525C strongly increases it [14]. Notably, double mutant cycle analyses of K538 and D411 mutants suggest these residues engage in a salt-bridge crucial for K_V_11.1 gating charge movement [40].

Using state-dependent cysteine accessibility assays, Zhang et al. showed that K525C is selectively accessible to extracellular MTSET only in depolarized states associated with inactivation, while remaining poorly accessible in the activated/open state and completely inaccessible in the closed state [45]. In contrast, cysteine substitutions at other S4 gating charges were never accessible to extracellular reagents, suggesting K525 occupies a unique position at the VSD extracellular boundary. Based on these measurements, the authors proposed that the closed state involves stabilizing two electrostatic interactions: K525–D456 and K538–D411 [14]. However, the structural identity of this “closed” state remains ambiguous; it is unclear if it represents the de-activated/closed state or the activated/closed state (described by the authors as open/inactivated). Cryo-EM structures of the EAG1 channel by Whicher and MacKinnon revealed a pore in a closed conformation despite a depolarized VSD, indicating pore closure can occur independently of full VSD deactivation [47]. Similarly, MD simulations of K_V_11.1 resting/closed state models [48] differ markedly from recently resolved activated/open high-K^+^ and activated/closed low-K^+^ K_V_11.1 structures [37], where the VSD remains activated (Fig. S7a) while the pore adopts conductive or non-conductive conformations (Fig. 3e-f).

These observations suggest that multiple non-conductive states exist in K_V_11.1, not all of which correspond to a fully deactivated VSD. Further insight comes from gating current measurements by Dou et al., who identified a fast component of gating charge movement (Q_fast_), implicating K538-D411 electrostatic interaction (Fig. 6b-d,Fig. S7a) in a local rearrangement of S4 that accompanies, rather than precedes, pore inactivation [40]. These studies additionally support the idea that S4 does not act solely as a voltage-dependent translocation element in K_V_11.1, but also participates in the conformational pathway that leads the channel toward its inactivated selectivity filter state. Perturbations of this pathway, such as those introduced by Class II missense mutations affecting S4 gating charges (including R534 ones) or its countercharges, could destabilize the energetic pathway toward inactivation, thereby preventing the channel from adopting conformations compatible with ER export. Indeed, R534C accelerates deactivation and induces depolarizing shifts in activation and inactivation voltage dependence without affecting inactivation kinetics [49]. Similarly, R534A shifts steady-state inactivation [13], and R534L demonstrates reduced surface expression and diminished ionic current in HEK293 cells [17]. Consistent with this view, variants such as D456Y [11], which may weaken the stabilizing electrostatic interaction with K525, exhibit trafficking deficiency, whereas charge-reversal missense mutations at R534 that preserve local electrostatic balance (Fig. 6b, right panel) can partially maintain this coupling and support trafficking through an anion-*π* interaction [50] with F463 in a transmembrane environment (Brian’s R534D unpublished results). Interestingly, three of the residues involved in our state-independent PD-C_LINKER_ bundle are associated with known LQT2 Class II missense mutations (T623I, R696C, R696P, P721L) and a Class III missense mutation (D767Y) [11, 33], while the F557L mutant was shown to impair the interaction of K_V_11.1 with pharmacological correctors [32, 41].

### 5.3 ’Correctability’ of trafficking-deficient missense variants

Some Class III missense mutations, such as Y652C, reportedly restore the trafficking of Class II mutants [19], which we attribute to the disruption the hydrophobic T623–Y652 interaction within the pore domain. This perturbation weakens the strong correlations between T623–Y652 and intersubunit C_LINKER_ interactions, which are characteristic of non-trafficking variants. As a consequence, Y652C perturbs the pore-domain coupling network, restoring the correlation patterns observed for T623–Y652 in Y652C, R531W–Y652C, and R534L–Y652C to values comparable to wild-type and other trafficking-competent variants. Based on these observations, K_V_11.1 blockers such as E-4031 or dofetilide [30, 31], which were reported to restore trafficking for a substantial subset of Class II missense mutations [33, 39, 44], including R534C [11] and R534L [17], may also restore the aforementioned correlation patterns as part of its trafficking correction mechanism. Notably, the high-affinity K_V_11.1 blocker E-4031 displayed strong state-dependence, preferentially binding to inactivated pore conformations [51, 52], whereas dofetilide bound K_V_11.1 in both open and inactivated conformations [52–55].

In contrast, many Class II variants located in the transmembrane region of K_V_11.1 remained uncorrectable, highlighting mechanistic heterogeneity in this channel’s ER export failure [33]. However, studies demonstrated that K_V_11.1 missense mutations characterized by an absence of inactivation and located on the conduction pathway, such as S620T [51] or S631A [53], trafficked normally while remaining insensitive to pharmacological correctors. This implies that K_V_11.1 inactivation ability is not a strict requirement for trafficking to the plasma membrane. Notably, many pharmacologically correctable Class II variants, such as R534A [13], R534C [49], and N488K [52], did not abolish K_V_11.1 inactivation but instead shifted its voltage-dependence without modifying its kinetics, suggesting that trafficking rescue requires preservation of access to an inactivation-linked conformational ensemble rather than intact inactivation kinetics per se. Lower temperatures (below 27 ^◦^C) also reportedly corrected trafficking-deficient K_V_11.1 mutants [33, 56]. This was explained by the lower temperature at which the functional studies were performed in oocytes (24-26 °C) compared to the temperature of the samples used for microscope imagery (37 ^◦^C). A similar observation was made for Western blots of the R534L mutant, which demonstrated decreased surface expression at 37 ^◦^C [17]. Moreover, multiple mutagenesis scanning of S4 was also performed at room temperature (below 27 ^◦^C), which might explain why R534 missense mutations still showed surface expression despite being associated with a Class II phenotype.

## 6 Conclusions

Together, these observations support the notion that trafficking defects in K_V_11.1 can arise from stabilization of non-native allosteric ensembles, whether initiated at the level of the voltage-sensing domain, the selectivity filter, or the C_LINKER_ itself. Specifically, these experimentally observed rescue patterns and our correlation analyses support the notion that K_V_11.1 trafficking is governed by the stability of a native allosteric ensemble, rather than individual local interactions or linear retention signals [57]. Furthermore, trafficking-deficient variants exhibit aberrant coupling between SF interactions and intersubunit contacts within the C_LINKER_, manifesting as an allosteric rigidification of the channel’s trans-membrane and cytoplasmic assembly. This contrasts with trafficking-competent variants, in which the SF and the C_LINKER_ remain dynamically decoupled, preserving the conformational flexibility required for maturation and ER export. Meanwhile, perturbations that stabilize these non-native coupling states, e.g. those arising from missense mutations in the VSD, the SF, or the C_LINKER_, promote recognition by the ER quality-control machinery and thereby defective trafficking. [11, 33].

## Supporting information

Supplementary Figures S1-S8 can be found in the Supplementary Material

## Glossary

RMSD: root mean squared deviations
RMSF: root mean squared fluctuations

## 7 Supplement

### S.1 Acknowledgements

Research reported in this publication was supported by the Maximizing Investigators’ Research Award (MIRA) (R35) from the National Institute of General Medical Sciences (NIGMS) of the National Institutes of Health (NIH) under grant number R35GM148284. American Heart Association under grant number 20IPA35320141. We thank Bin Sun for helpful discussions and for establishing the simulation protocol that guided subsequent simulations. This work used the Extreme Science and Engineering Discovery Environment (XSEDE) [**Towns2014**], which is supported by National Science Foundation grant number ACI-1548562.

### S.2 Supp Results

**Figure S1:**
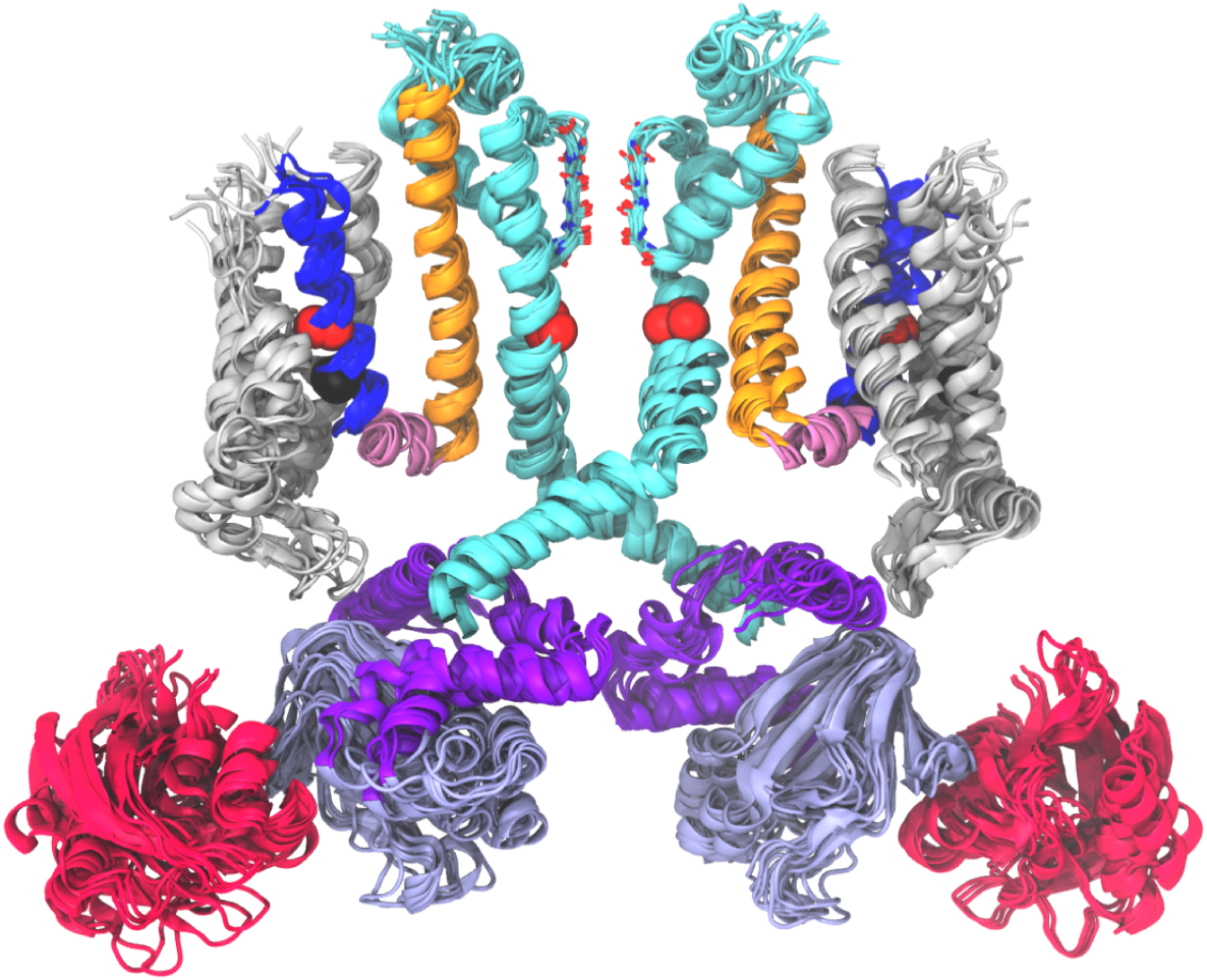
Structural alignment of all hERG variant models. For each variant, two monomers are shown for clarity. The cartoon representation of hERG shows the PAS domain in pink, the VSD in gray, S4 in blue, S5 in orange, and the rest of the pore domain in cyan. The CNBD is depicted in lilac. The positions of the mutated residues are shown in spheres.

**Figure S2:**
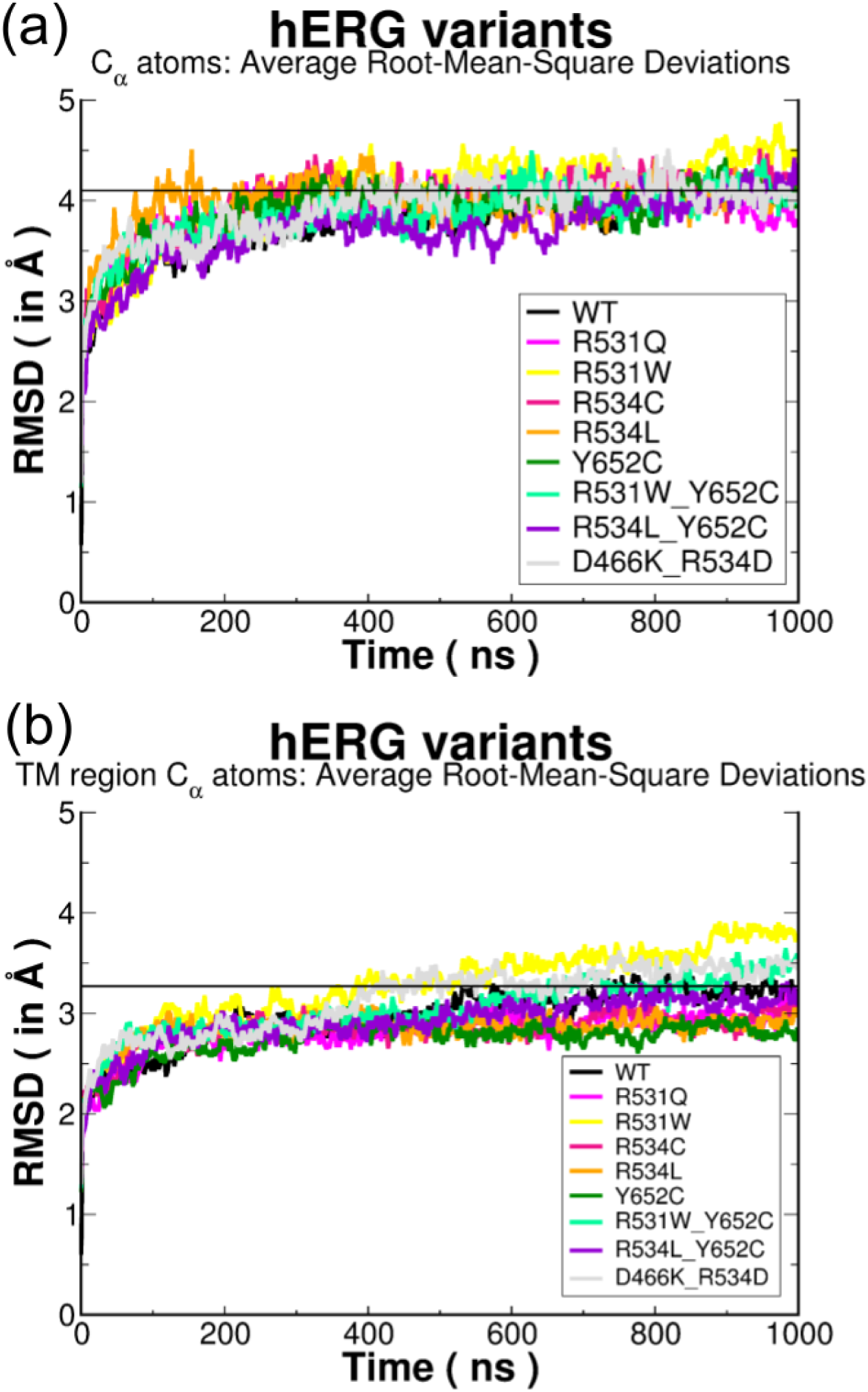
RMSD calculations of hERG MD trajectories. Average C*_α_* atom RMSD of the whole channel (a) and transmembrane domains (b) from the 3 × 1000 *µ*s MD simulations of hERG variants.

**Figure S3:**
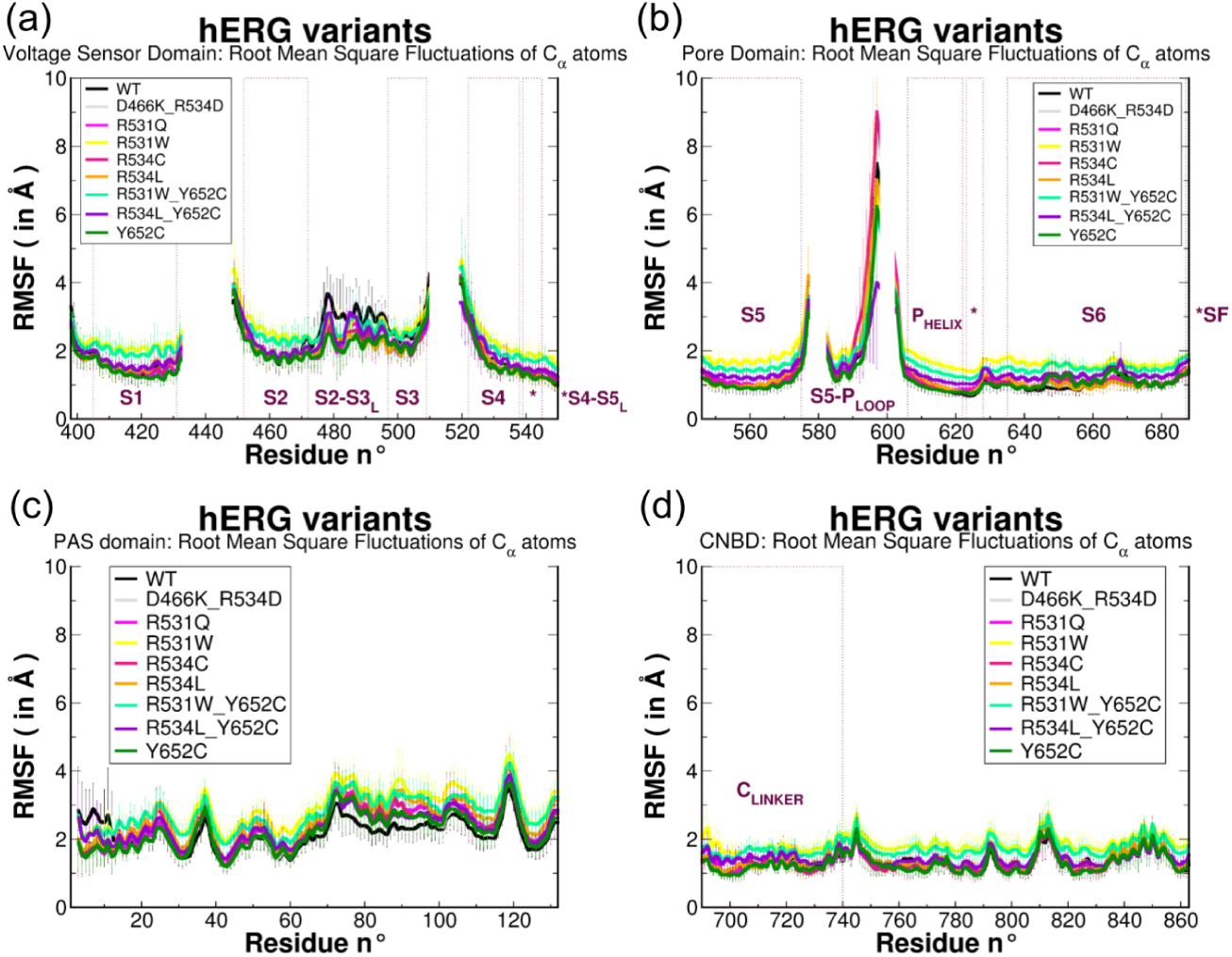
RMSF calculations of hERG MD trajectories. Average C*_α_* atom RMSF of the voltage sensor domain (a), the pore domain (b), the PAS domain (c) and the CNBD (d) from the 3 × 1000 *µ*s MD simulations of hERG variants.

**Figure S4:**
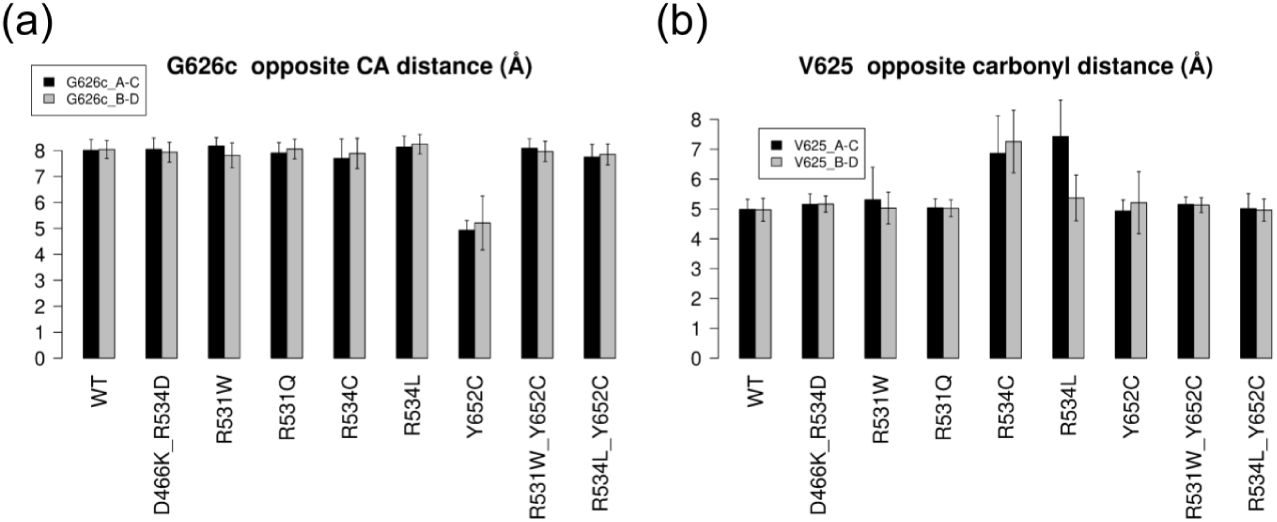
Key average opposite subunit distances in the selectivity filter in the MD trajectories of WT hERG and its variants. (a) Bar plot showing the average distance between the V625 backbone oxygen atoms of opposing subunits. (b) Bar plot showing the average distance between the G626 C*_α_* atoms of opposing subunits (labeled A, B, C, D).

**Figure S5:**
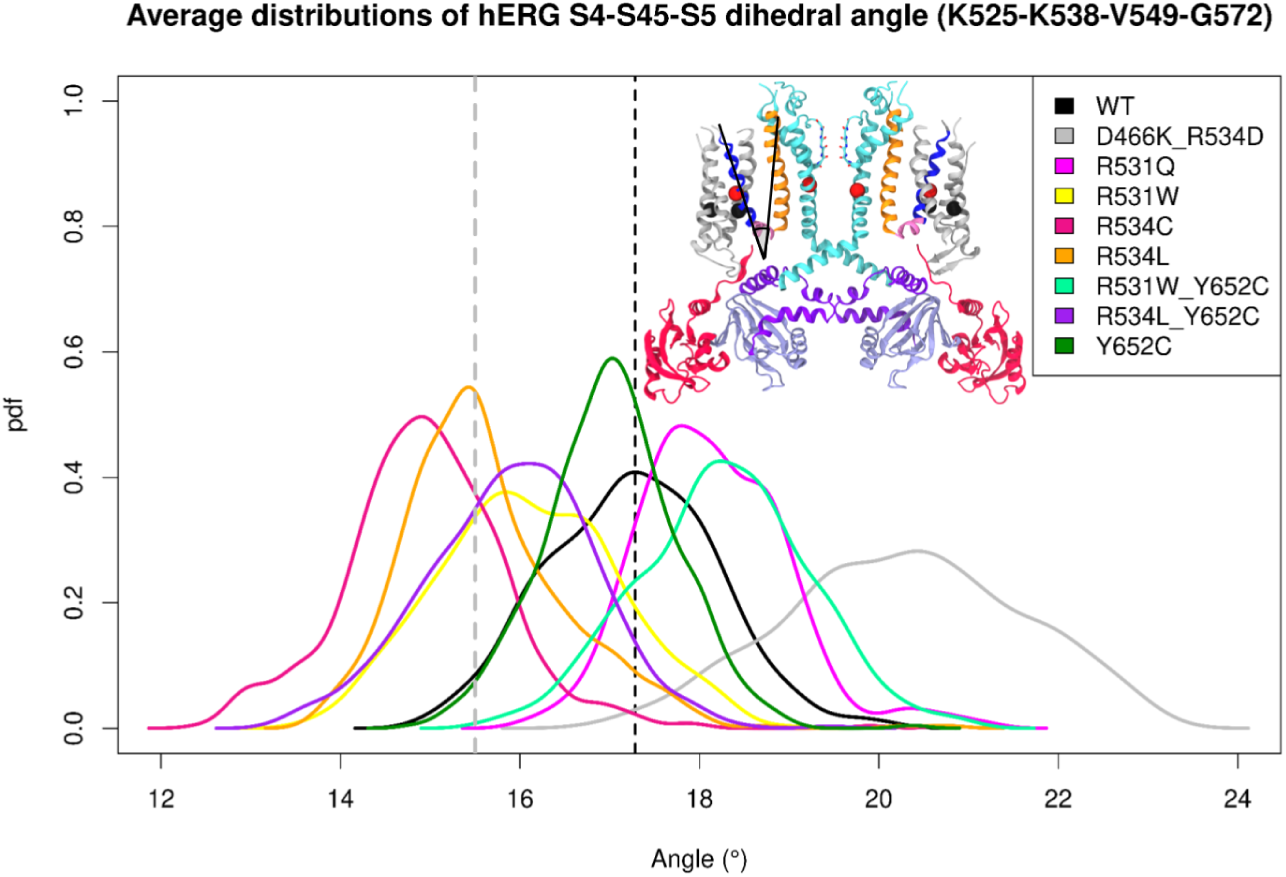
Probability densities of the the time-dependent torsion angle between the S4 and S5 helices relative to the S4-S5*_L_* axis. through-out the MD simulation of hERG WT and variant models. The most frequent angle value of WT hERG and the threshold value between hERG trafficking-competent and non-trafficking variants most frequent values are represented by black and gray vertical dashed lines, respectively.

**Figure S6:**
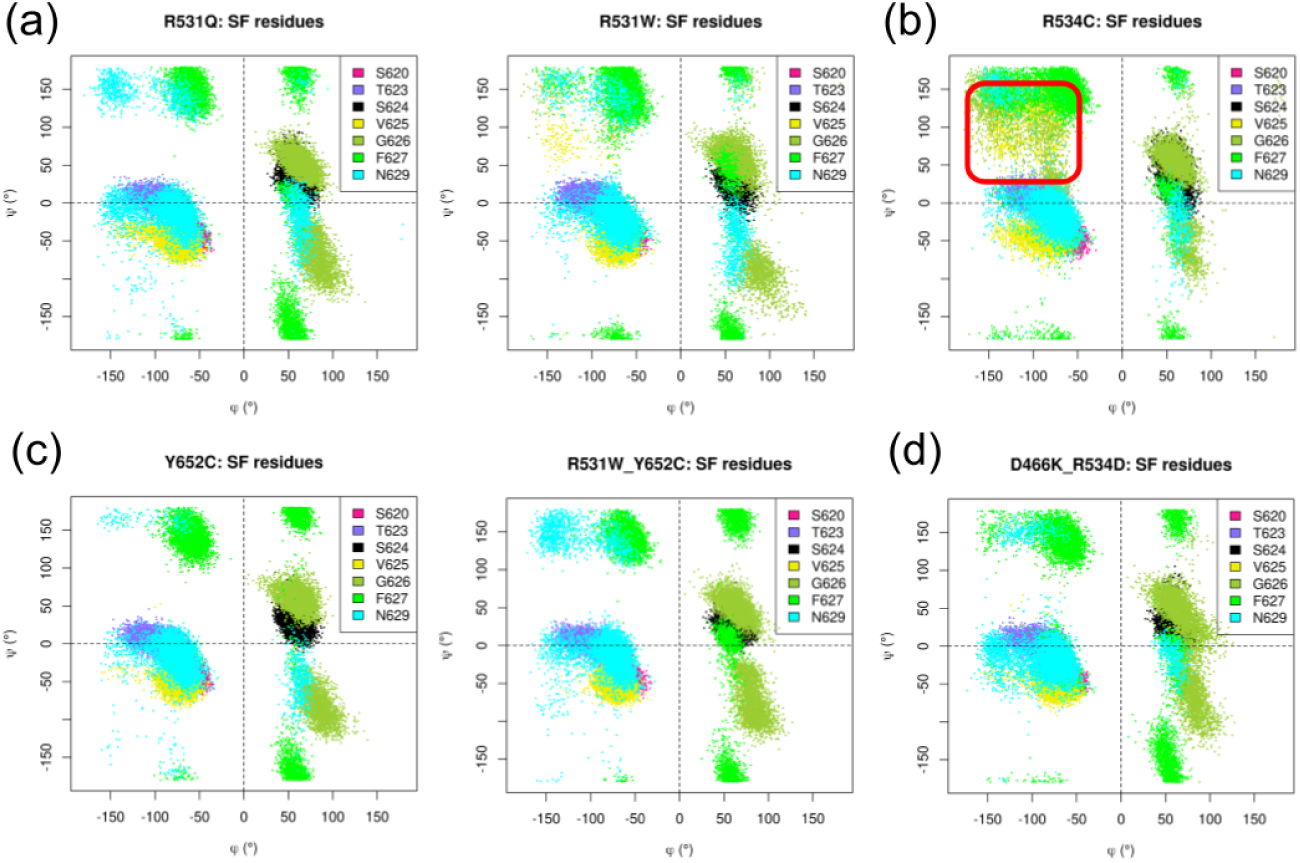
Ramachandran plots of SF residues in hERG Class II and Class III variants. The Ramachandran plots display the *φ* and *ψ* torsion angles of hERG residues S620, T623, S624, V625, G626, F627 and N629, all throughout the MD trajectories of (a) Non-conductive mutants R531Q (left) and R531W (right), (b) the non-trafficking mutant R534C, (c) Class III Y652C single mutant (left) and R531W-Y652C double mutant (right), and (d) the trafficking-corrected double mutant D466K-R534D from the 3 × 1000 *µ*s MD simulations of hERG variants. Interaction probabilities were estimated by integrating distance probability densities below a set of physically motivated thresholds, as described in detail in previous studies [58, 59]. These interaction probabilities reflect the likelihood that geometries compatible with a given interaction are sampled during the MD trajectories. All these time-dependent calculations were performed with the use of scripts written in the Tcl language that is implemented within VMD program [60, 61]. Protein-protein interactions were quantified by measuring specific inter-atomic distances, Interaction probabilities were then quantified by integrating the probability density function of the corresponding interatomic distance over a distance range consistent with the interaction type (salt-bridges and hydrogen bonds: 3.5 °A; hydrophobic contacts, *π*-*π* stacking, cation-*π*, anion-*π* and CH-*π*: 5 °A). As an example, salt-bridge / H-bond probabilities are reported here as integrated probability densities, computed as Riemann sums of the corresponding interatomic distance distributions below 3.5 °A. Sulfur-*π* and sulfur-aliphatic contacts, due to their significant binding energies in transmembrane proteins [62] were also considered, using a threshold of 6 °A for Met-*π* interactions, and 5 °A for Cys-*π* and Sulfur-aliphatic interactions.

**Figure S7:**
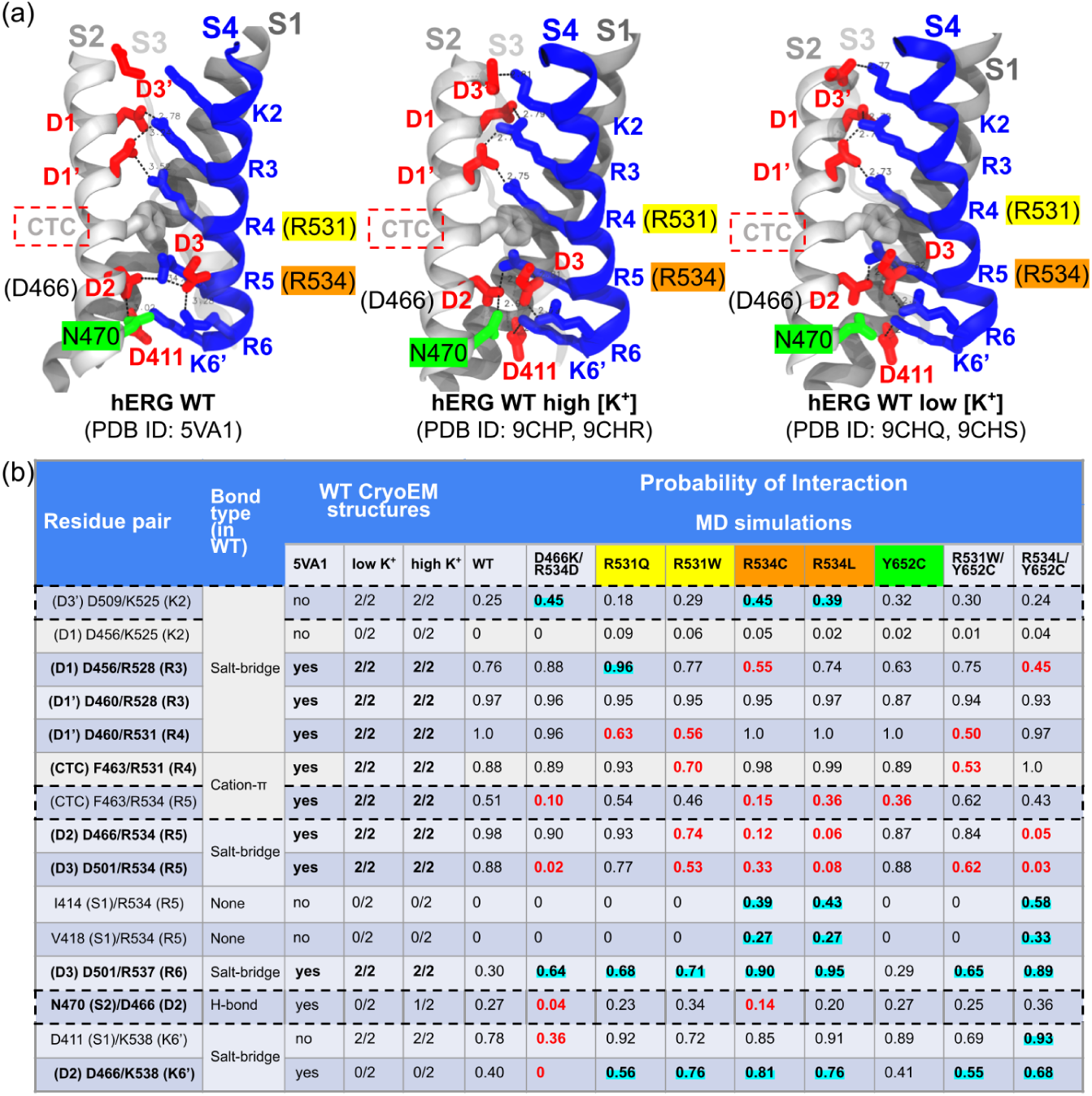
Identifying K_V_11.1 trafficking-dependent interactions in the pore. (a) Structural mapping of conserved S2 (solid gray), S3 (transparent gray), and S4 (blue) residues and interactions in hERG experimental models: The first Cryo-EM structure (left), and more recent hERG Cryo-EM structures, each resolved in high (middle) and low (right) [K^+^]_out_ concentrations. Residues are shown as sticks; acidic residues are depicted in red, basic residues in blue. The CTC is colored in gray. (b) Interaction probabilities of S4 electrostatic interactions throughout MD trajectories of wild-type K_V_11.1 and variants. Values highlighted in red are at least 0.14 lower than WT, while values in cyan are 0.14 higher than WT. Residue pairs present in the K_V_11.1 Cryo-EM structure (PDB: 5VA1 [20]) are bolded.

**Figure S8:**
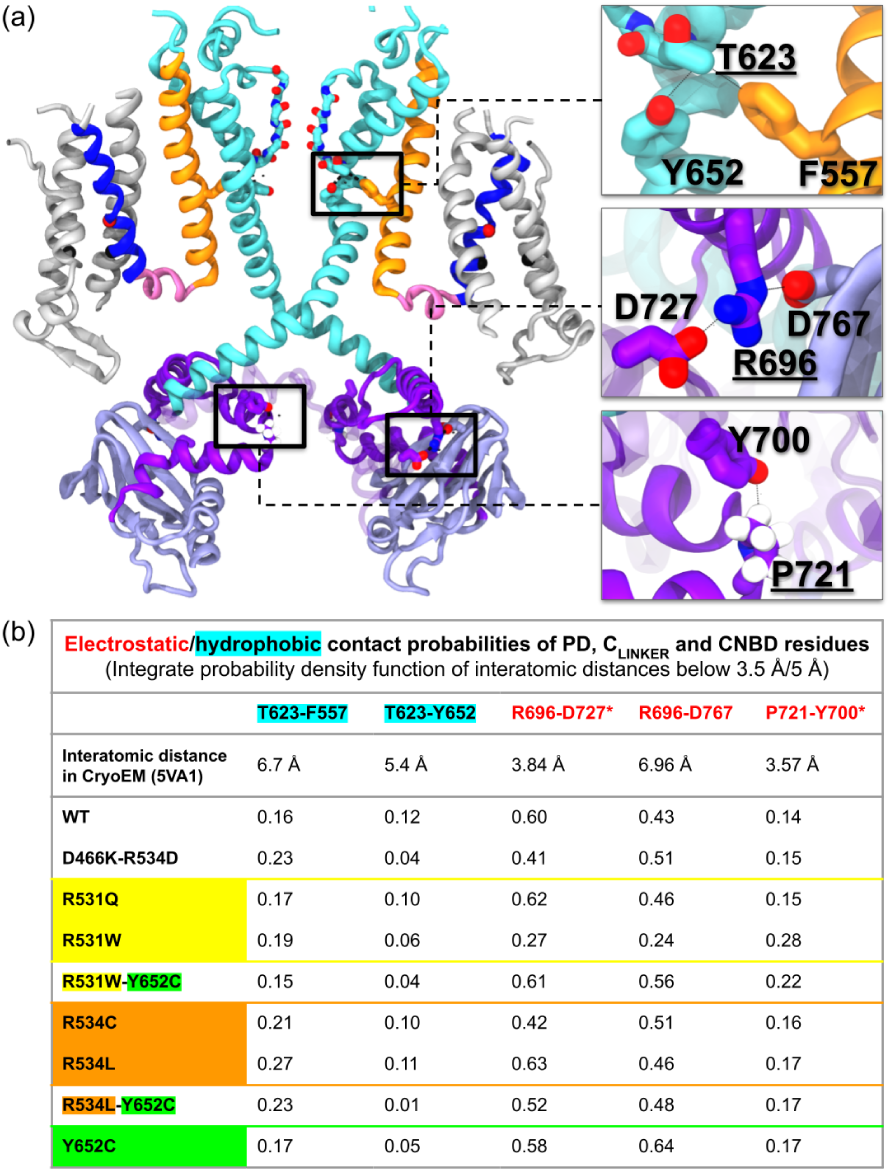
K_V_11.1 PD-C_LINKER_ coupling network: variant-independent interactions. (a) The left panel shows a cartoon representation of two subunits of K_V_11.1’s TM region (S1-S3 in grey, S4 in blue, S4-S5_L_ in pink, S5 in orange, and S5-P loop, P_HELIX_ and S6 in cyan), along with the C_LINKER_ (purple) and the CNBD (lilac) of two adjacent subunits. A third C_LINKER_, depicted in transparent purple, is adjacent to the cytosolic subunits depicted in solid colors and belongs to the TM subunit on the right. The residues involved in variant-independent interactions are framed in black, and their extended views of their stick representations are shown on the right panel. The residues that are associated with Class II LQT2 missense mutations have underlined labels on the right panel. (b) The tab reports the interactions probabilities of the residues depicted in (a). Intersubunit residue pairs are marked with an asterisk. To calculate the integrated distance density functions, thresholds of 3.5 and 5 °A were used for the residue pairs labeled in red and cyan, respectively. A selection of the aforementioned interatomic distances were also used to generate correlation matrices that report Pearson coefficients, each estimated at a p-value of 0.05, using the ggcorrplot package [63] in scripts written in R language. Correlation matrices were constructed from time-resolved descriptors to capture allosteric coupling.

**Figure S9:**
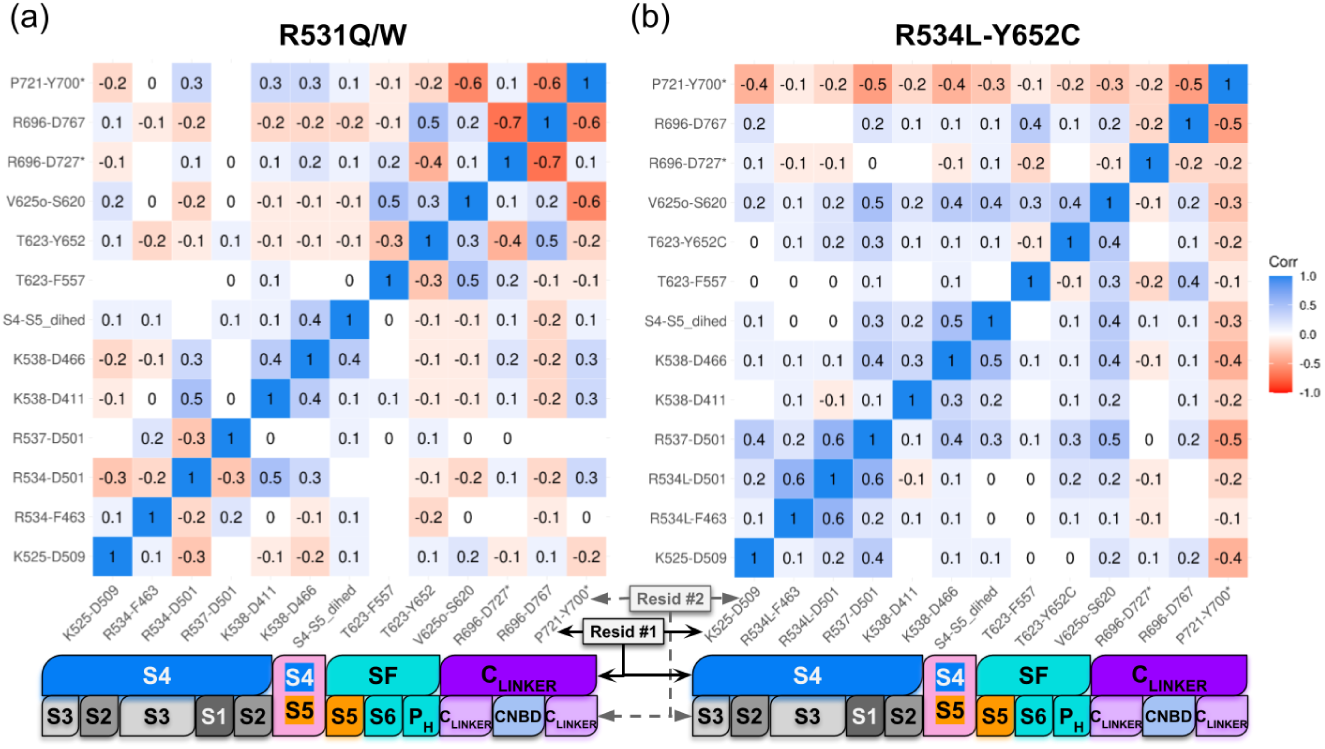
Correlation matrices of pore domain interhelical interactions in Kv11.1 non-conducting R531 variants and trafficking-correcting variant. (a) Pearson coefficient heatmap of interatomic distances of pore domain residue pairs in MD trajectories of (a) R531Q and R531W Kv11.1 and (b) the trafficking corrected double mutant R534L-Y652C. For each residue pair, the Pearson coefficient was calculated with p ¡ 0.05; blank areas represent non-significant values. In both panels, two horizontal bars are colored according to the region of each residue in the corresponding residue pairs.

**Figure S10:**
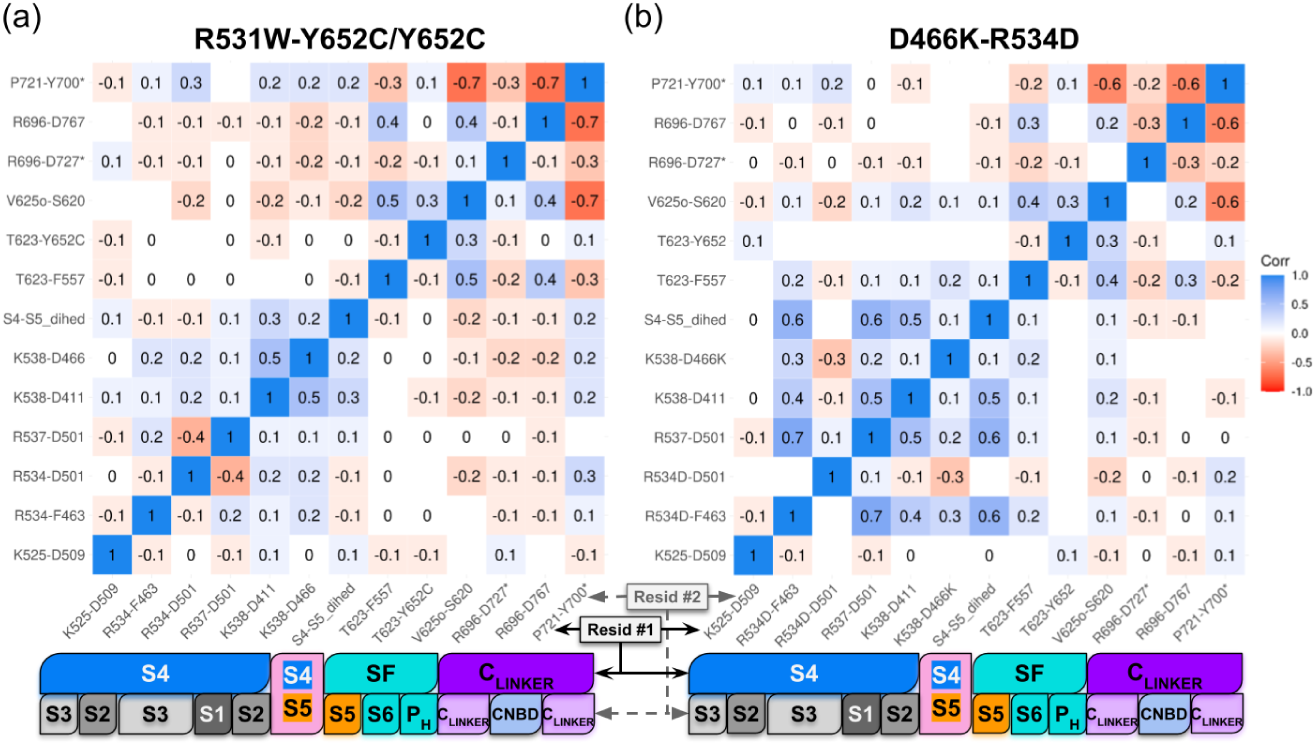
Correlation matrices of pore domain residue interhelical interactions in hERG non-conducting Y652C single and double mutants and charge-reversal mutant D466K-R534D. (a) Pearson correlation coefficient heatmap of the interatomic distances of pore domain residue pairs in the MD trajectories of (a) Y652C and R531W-Y652C hERG and (b) hERG variant D466K-R534D. For each couple of residue pairs, the Pearson coefficient was calculated with a p-value of 0.05, and blank areas represent non significant values. In both panels, two horizontal bars are colored according to the region of each residue in the corresponding residue pairs.

