## Supplementary Figures S1-S8 can be found in the Supplementary Material for "Allosteric Mechanisms Underlying Long QT Syndrome Type 2 (LQT2)-Associated Mutations in hERG Channels"

April 2, 2026

### 7 Supplement

#### S.1 Acknowledgements

Research reported in this publication was supported by the Maximizing Investigators' Research Award (MIRA) (R35) from the National Institute of General Medical Sciences (NIGMS) of the National Institutes of Health (NIH) under grant number R35GM148284. American Heart Association under grant number 20IPA35320141. We thank Bin Sun for helpful discussions and for establishing the simulation protocol that guided subsequent simulations. This work used the Extreme Science and Engineering Discovery Environment (XSEDE) [**Towns2014**], which is supported by National Science Foundation grant number ACI-1548562.

#### S.2 Supp Results

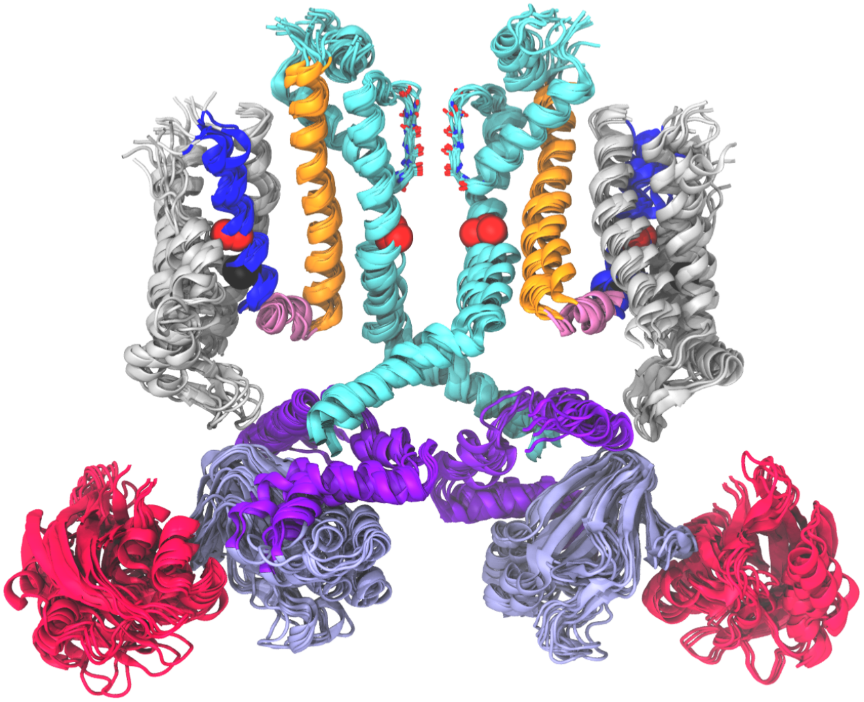

Figure S1: **Structural alignment of all hERG variant models.** For each variant, two monomers are shown for clarity. The cartoon representation of hERG shows the PAS domain in pink, the VSD in gray, S4 in blue, S5 in orange, and the rest of the pore domain in cyan. The CNBD is depicted in lilac. The positions of the mutated residues are shown in spheres..

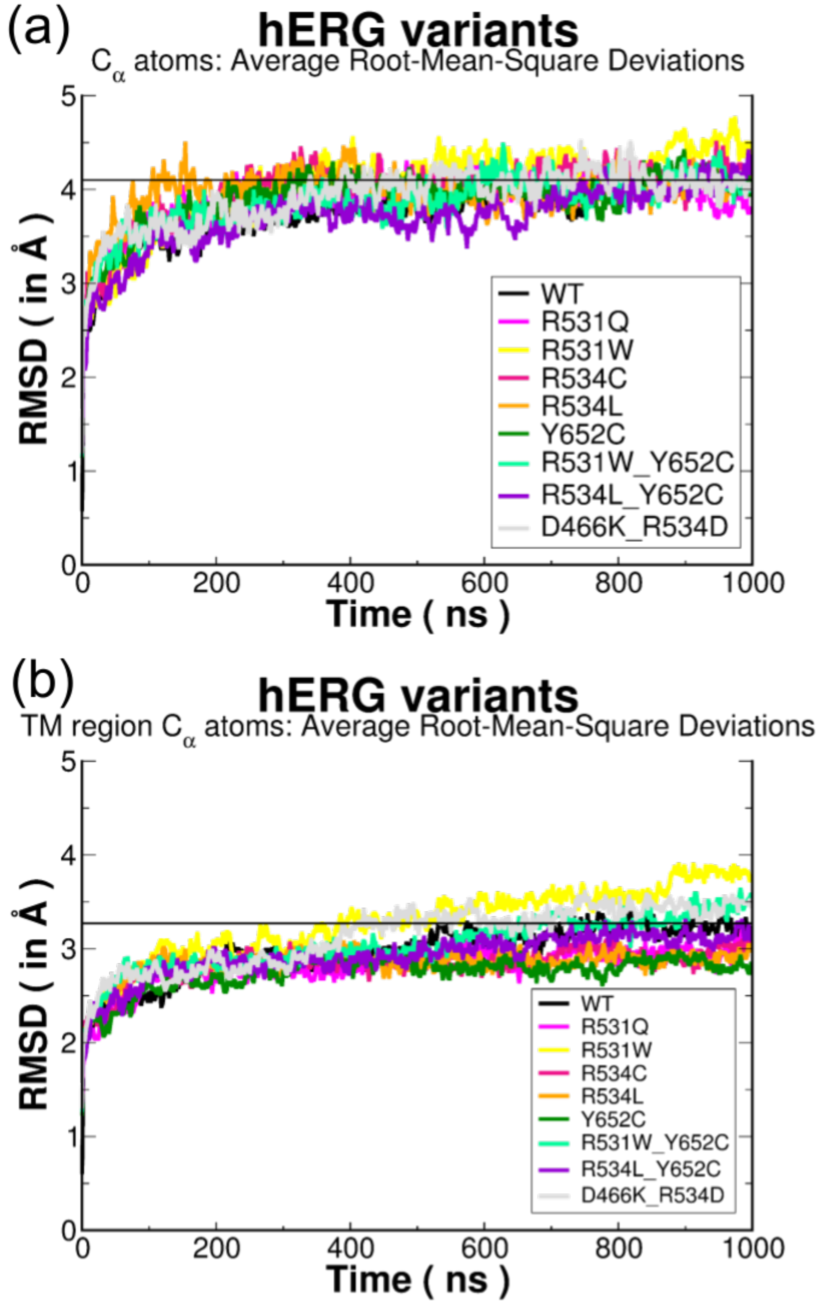


Figure S2: **RMSD calculations of hERG MD trajectories.** Average $C_\alpha$ atom RMSD of the whole channel (a) and transmembrane domains (b) from the $3 \times 1000$ $\mu$s MD simulations of hERG variants.

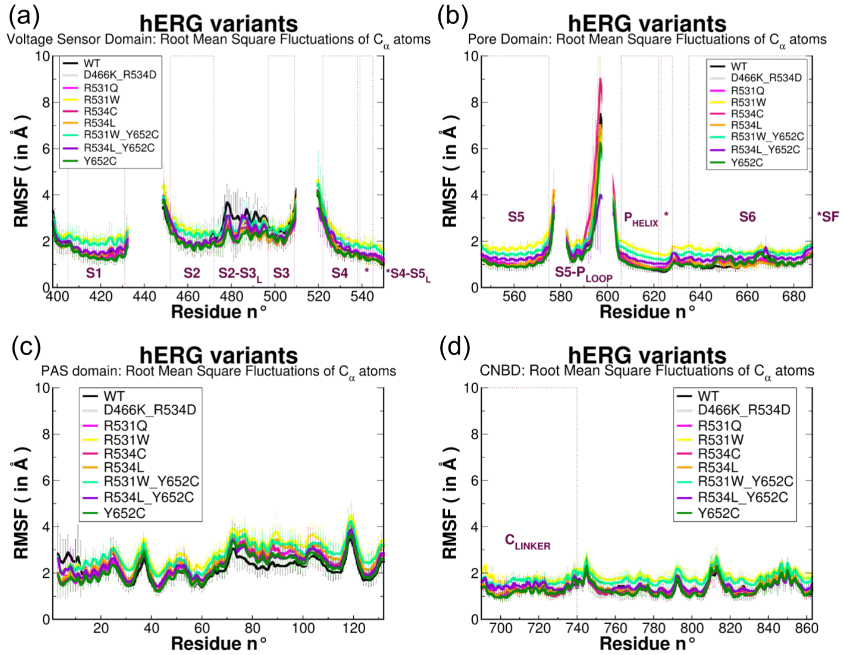


Figure S3: **RMSF calculations of hERG MD trajectories.** Average $C_\alpha$ atom RMSF of the voltage sensor domain (a), the pore domain (b), the PAS domain (c) and the CNBD (d) from the 3 × 1000 $\mu$s MD simulations of hERG variants.

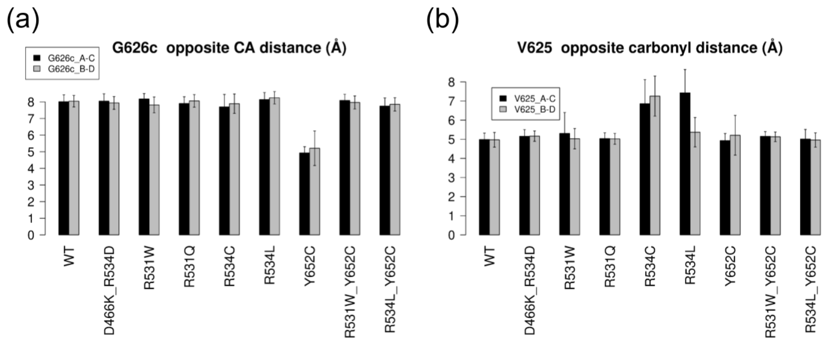


Figure S4: **Key average opposite subunit distances in the selectivity filter in the MD trajectories of WT hERG and its variants.** (a) Bar plot showing the average distance between the V625 backbone oxygen atoms of opposing subunits. (b) Bar plot showing the average distance between the G626 $C_\alpha$ atoms of opposing subunits (labeled A, B, C, D).

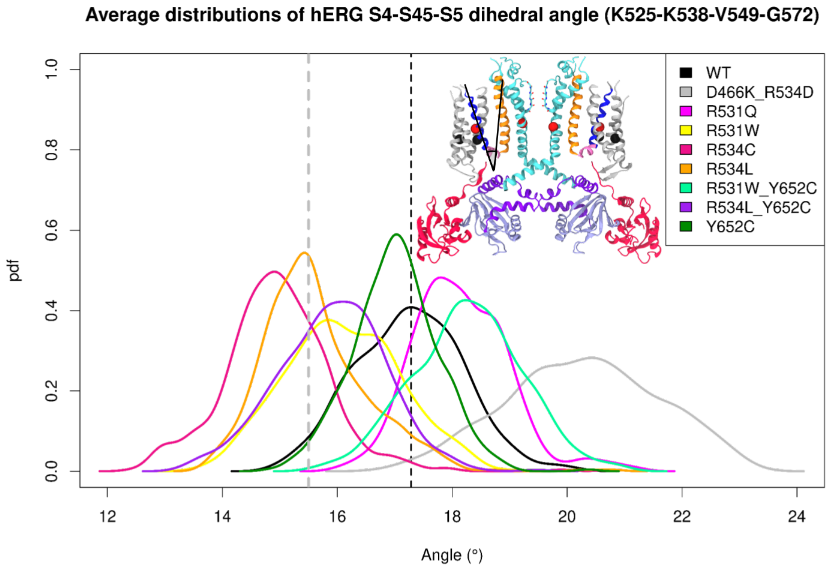


Figure S5: **Probability densities of the the time-dependent torsion angle between the S4 and S5 helices relative to the S4-S5$_L$ axis** throughout the MD simulation of hERG WT and variant models. The most frequent angle value of WT hERG and the threshold value between hERG trafficking-competent and non-trafficking variants most frequent values are represented by black and gray vertical dashed lines, respectively. .

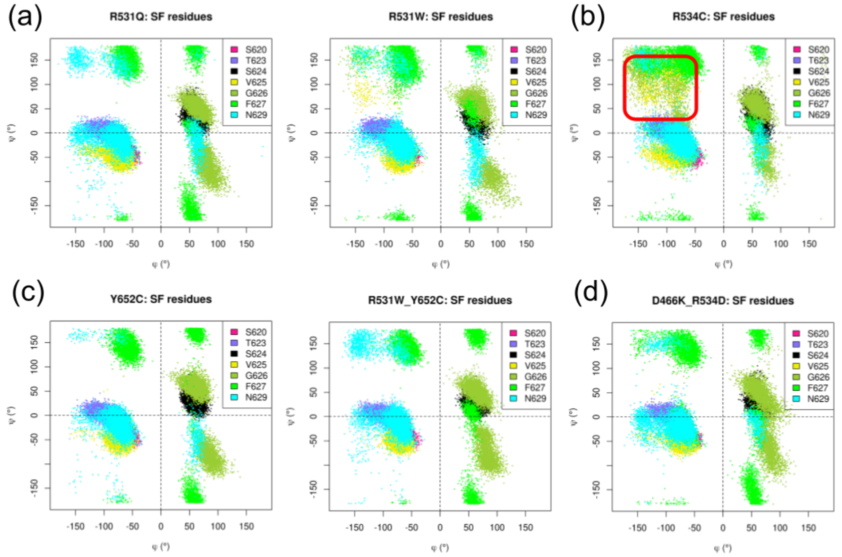


Figure S6: **Ramachandran plots of SF residues in hERG Class II and Class III variants** The Ramachandran plots display the $\phi$ and $\psi$ torsion angles of hERG residues S620, T623, S624, V625, G626, F627 and N629, all throughout the MD trajectories of (a) Non-conductive mutants R531Q (left) and R531W (right), (b) the non-trafficking mutant R534C, (c) Class III Y652C single mutant (left) and R531W-Y652C double mutant (right), and (d) the trafficking-corrected double mutant D466K-R534D from the 3 $\times$ 1000 $\mu$s MD simulations of hERG variants.

Interaction probabilities were estimated by integrating distance probability densities below a set of physically motivated thresholds, as described in detail in previous studies [58, 59]. These interaction probabilities reflect the likelihood that geometries compatible with a given interaction are sampled during the MD trajectories. All these time-dependent calculations were performed with the use of scripts written in the Tcl language that is implemented within VMD program [60, 61]. Protein-protein interactions were quantified by measuring specific interatomic distances, Interaction probabilities were then quantified by integrating the probability density function of the corresponding interatomic distance over a distance range consistent with the interaction type (salt-bridges and hydrogen bonds: 3.5 Å; hydrophobic contacts, $\pi$-$\pi$ stacking, cation-$\pi$, anion-$\pi$ and CH-$\pi$: 5 Å). As an example, salt-bridge / H-bond probabilities are reported here as integrated probability densities, computed as Riemann sums of the corresponding interatomic distance distributions below 3.5 Å. Sulfur-$\pi$ and sulfur-aliphatic contacts, due to their significant binding energies in transmembrane proteins [62] were also considered, using a threshold of 6 Å for Met-$\pi$ interactions, and 5 Å for Cys-$\pi$ and Sulfur-aliphatic interactions.

(a)

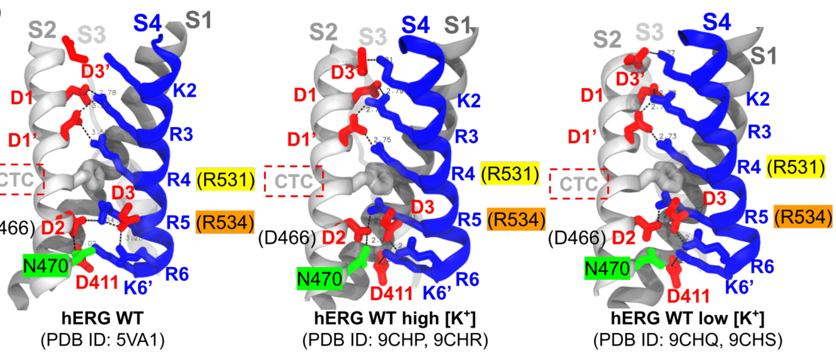


(b)

| Residue pair | Bond type (in WT) | WT CryoEM structures | | | Probability of Interaction MD simulations | | | | | | | | |
|---|---|---|---|---|---|---|---|---|---|---|---|---|---|
| | | 5VA1 | low K+ | high K+ | WT | D466K/R534D | R531Q | R531W | R534C | R534L | Y652C | R531W/Y652C | R534L/Y652C |
| (D3') D509/K525 (K2) | Salt-bridge | no | 2/2 | 2/2 | 0.25 | 0.45 | 0.18 | 0.29 | 0.45 | 0.39 | 0.32 | 0.30 | 0.24 |
| (D1) D456/K525 (K2) | | no | 0/2 | 0/2 | 0 | 0 | 0.09 | 0.06 | 0.05 | 0.02 | 0.02 | 0.01 | 0.04 |
| **(D1) D456/R528 (R3)** | | **yes** | **2/2** | **2/2** | 0.76 | 0.88 | 0.96 | 0.77 | 0.55 | 0.74 | 0.63 | 0.75 | 0.45 |
| **(D1') D460/R528 (R3)** | | **yes** | **2/2** | **2/2** | 0.97 | 0.96 | 0.95 | 0.95 | 0.95 | 0.97 | 0.87 | 0.94 | 0.93 |
| **(D1') D460/R531 (R4)** | | **yes** | **2/2** | **2/2** | 1.0 | 0.96 | 0.63 | 0.56 | 1.0 | 1.0 | 1.0 | 0.50 | 0.97 |
| **(CTC) F463/R531 (R4)** | Cation-π | **yes** | **2/2** | **2/2** | 0.88 | 0.89 | 0.93 | 0.70 | 0.98 | 0.99 | 0.89 | 0.53 | 1.0 |
| (CTC) F463/R534 (R5) | | **yes** | **2/2** | **2/2** | 0.51 | 0.10 | 0.54 | 0.46 | 0.15 | 0.36 | 0.36 | 0.62 | 0.43 |
| **(D2) D466/R534 (R5)** | Salt-bridge | **yes** | **2/2** | **2/2** | 0.98 | 0.90 | 0.93 | 0.74 | 0.12 | 0.06 | 0.87 | 0.84 | 0.05 |
| **(D3) D501/R534 (R5)** | | **yes** | **2/2** | **2/2** | 0.88 | 0.02 | 0.77 | 0.53 | 0.33 | 0.08 | 0.88 | 0.62 | 0.03 |
| I414 (S1)/R534 (R5) | None | no | 0/2 | 0/2 | 0 | 0 | 0 | 0 | 0.39 | 0.43 | 0 | 0 | 0.58 |
| V418 (S1)/R534 (R5) | None | no | 0/2 | 0/2 | 0 | 0 | 0 | 0 | 0.27 | 0.27 | 0 | 0 | 0.33 |
| **(D3) D501/R537 (R6)** | Salt-bridge | **yes** | **2/2** | **2/2** | 0.30 | 0.64 | 0.68 | 0.71 | 0.90 | 0.95 | 0.29 | 0.65 | 0.89 |
| **N470 (S2)/D466 (D2)** | H-bond | yes | 0/2 | 1/2 | 0.27 | 0.04 | 0.23 | 0.34 | 0.14 | 0.20 | 0.27 | 0.25 | 0.36 |
| D411 (S1)/K538 (K6') | Salt-bridge | no | 2/2 | 2/2 | 0.78 | 0.36 | 0.92 | 0.72 | 0.85 | 0.91 | 0.89 | 0.69 | 0.93 |
| **(D2) D466/K538 (K6')** | | yes | 0/2 | 0/2 | 0.40 | 0 | 0.56 | 0.76 | 0.81 | 0.76 | 0.41 | 0.55 | 0.68 |

Figure S7: **Identifying $K_V$11.1 trafficking-dependent interactions in the pore** (a) Structural mapping of conserved S2 (solid gray), S3 (transparent gray), and S4 (blue) residues and interactions in hERG experimental models: The first Cryo-EM structure (left), and more recent hERG Cryo-EM structures, each resolved in high (middle) and low (right) $[K^+]_{out}$ concentrations. Residues are shown as sticks; acidic residues are depicted in red, basic residues in blue. The CTC is colored in gray. (b) Interaction probabilities of S4 electrostatic interactions throughout MD trajectories of wild-type $K_V$11.1 and variants. Values highlighted in red are at least 0.14 lower than WT, while values in cyan are 0.14 higher than WT. Residue pairs present in the $K_V$11.1 Cryo-EM structure (PDB: 5VA1 [20]) are bolded. .

(a)

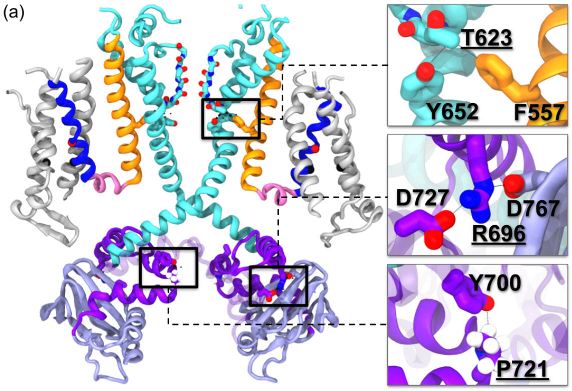


(b)

| Electrostatic/hydrophobic contact probabilities of PD, $C_{LINKER}$ and CNBD residues (Integrate probability density function of interatomic distances below 3.5 Å/5 Å) | | | | | |
|---|---|---|---|---|---|
| | T623-F557 | T623-Y652 | R696-D727* | R696-D767 | P721-Y700* |
| Interatomic distance in CryoEM (5VA1) | 6.7 Å | 5.4 Å | 3.84 Å | 6.96 Å | 3.57 Å |
| WT | 0.16 | 0.12 | 0.60 | 0.43 | 0.14 |
| D466K-R534D | 0.23 | 0.04 | 0.41 | 0.51 | 0.15 |
| R531Q | 0.17 | 0.10 | 0.62 | 0.46 | 0.15 |
| R531W | 0.19 | 0.06 | 0.27 | 0.24 | 0.28 |
| R531W-Y652C | 0.15 | 0.04 | 0.61 | 0.56 | 0.22 |
| R534C | 0.21 | 0.10 | 0.42 | 0.51 | 0.16 |
| R534L | 0.27 | 0.11 | 0.63 | 0.46 | 0.17 |
| R534L-Y652C | 0.23 | 0.01 | 0.52 | 0.48 | 0.17 |
| Y652C | 0.17 | 0.05 | 0.58 | 0.64 | 0.17 |

Figure S8: **$K_V$11.1 PD-$C_{LINKER}$ coupling network: variant-independent interactions** (a) The left panel shows a cartoon representation of two subunits of $K_V$11.1's TM region (S1-S3 in grey, S4 in blue, S4-$S5_L$ in pink, S5 in orange, and S5-P loop, $P_{HELIX}$ and S6 in cyan), along with the $C_{LINKER}$ (purple) and the CNBD (lilac) of two adjacent subunits. A third $C_{LINKER}$, depicted in transparent purple, is adjacent to the cytosolic subunits depicted in solid colors and belongs to the TM subunit on the right. The residues involved in variant-independent interactions are framed in black, and their extended views of their stick representations are shown on the right panel. The residues that are associated with Class II LQT2 missense mutations have underlined labels on the right panel. (b) The tab reports the interactions probabilities of the residues depicted in (a). Intersubunit residue pairs are marked with an asterisk. To calculate the integrated distance density functions, thresholds of 3.5 and 5 Å were used for the residue pairs labeled in red and cyan, respectively. .

A selection of the aforementioned interatomic distances were also used to generate correlation matrices that report Pearson coefficients, each estimated at a p-value of 0.05, using the ggcorrplot package [63] in scripts written in R language. Correlation matrices were constructed from time-resolved descriptors to capture allosteric coupling.

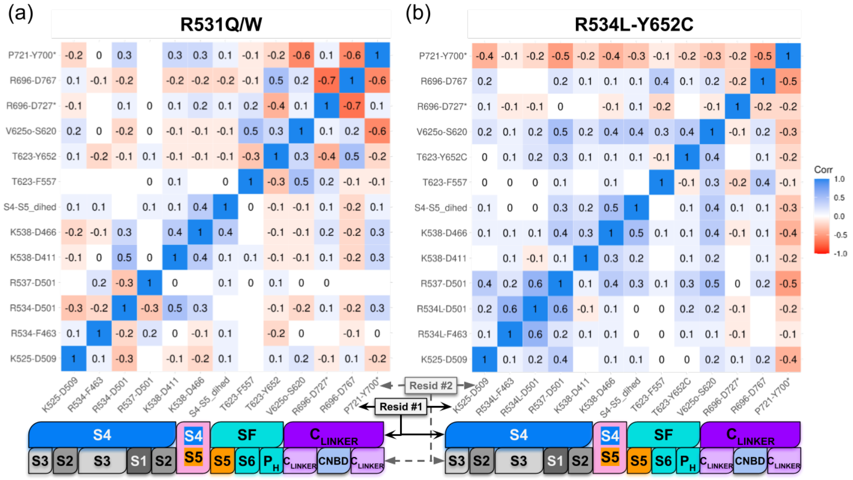


Figure S9: **Correlation matrices of pore domain interhelical interactions in Kv11.1 non-conducting R531 variants and trafficking-correcting variant** (a) Pearson coefficient heatmap of interatomic distances of pore domain residue pairs in MD trajectories of (a) R531Q and R531W Kv11.1 and (b) the trafficking corrected double mutant R534L-Y652C. For each residue pair, the Pearson coefficient was calculated with p ¡ 0.05; blank areas represent non-significant values. In both panels, two horizontal bars are colored according to the region of each residue in the corresponding residue pairs. .

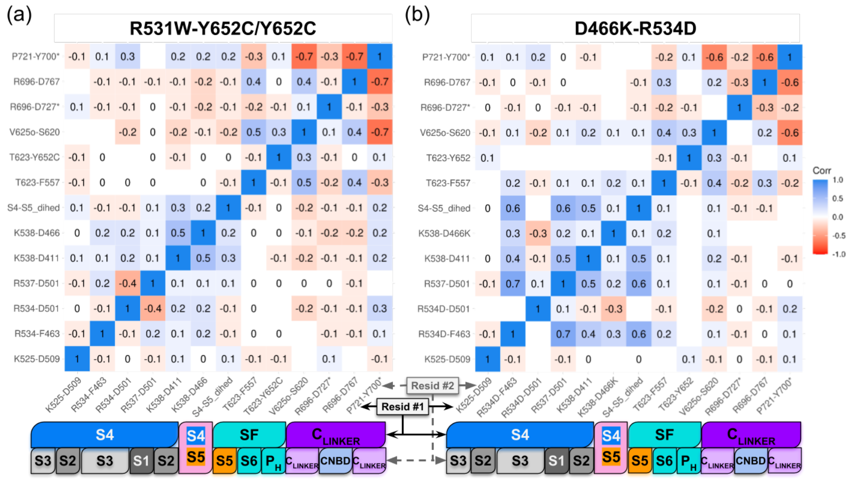


Figure S10: **Correlation matrices of pore domain residue interhelical interactions in hERG non-conducting Y652C single and double mutants and charge-reversal mutant D466K-R534D** (a) Pearson correlation coefficient heatmap of the interatomic distances of pore domain residue pairs in the MD trajectories of (a) Y652C and R531W-Y652C hERG and (b) hERG variant D466K-R534D. For each couple of residue pairs, the Pearson coefficient was calculated with a p-value of 0.05, and blank areas represent non significant values. In both panels, two horizontal bars are colored according to the region of each residue in the corresponding residue pairs. .
